# Unequal genetic redundancies among MYC bHLH transcription factors underlie seedling photomorphogenesis in Arabidopsis

**DOI:** 10.1101/2024.05.07.592999

**Authors:** Vikas Garhwal, Sreya Das, Sreeramaiah N. Gangappa

## Abstract

Light is one of the most critical ecological cues controlling plant growth and development. Plants have evolved complex mechanisms to cope with fluctuating light signals. In Arabidopsis, bHLH transcription factors MYC2, MYC3, and MYC4 have been shown to play a vital role in protecting plants against herbivory and necrotrophic pathogens. While the role of MYC2 in light-mediated seedling development has been studied in some detail, the role of MYC3 and MYC4 still needs to be discovered. Here, we show that MYC4 negatively regulates seedling photomorphogenesis, while the MYC3 function seems redundant. However, the genetic analysis reveals that MYC3/MYC4 together act as positive regulators of seedling photomorphogenic growth as the *myc3myc4* double mutants showed exaggerated hypocotyl growth compared to *myc4* single mutants and Col-0. Intriguingly, the loss of *MYC2* function in the *myc3myc4* double mutant background (*myc2myc3myc4*) resulted in further enhancement in the hypocotyl growth than *myc3myc4* double mutants in WL, BL and FRL, suggesting that MYC2/3/4 together play an essential and positive role in meditating optimal seedling photomorphogenesis. Besides, MYC3/MYC4 genetically and physically interact with HY5 to partially inhibit its function in controlling hypocotyl and photo-pigment accumulation. Moreover, our results suggest that COP1 physically interacts and degrades MYC3 and MYC4 through the 26S proteasomal pathway and controls their response to dark and light for fine-tuning HY5 function and seedling photomorphogenesis.

## Introduction

Light is a powerful stimulus that regulates plant development and phenotypic plasticity. Plant development, including seed germination, photomorphogenesis, leaf development, petiole elongation, shade avoidance, and flowering, is dependent on the presence or absence of light (Kendrick and Kronenberg, 1994; Wang and Deng, 2003; Jiao et al., 2007)(Cheng *et al*., 2021, Krahmer and Fankhauser, 2023, Huq *et al*., 2024). Light drives transition in the development of plants: Photomorphogenesis or de-etiolation happens as soon as a seedling emerges from the soil and sees the light for the first time. There are two contrasting patterns in the seedlings stage: Skotomorphogenesis and Photomorphogenesis. Several physiological responses change with the transition from skotomorphogenic to photomorphogenic seedling development, like opening of apical hook, expansion of cotyledons, inhibition of hypocotyl elongation, far-red light-controlled blocking of greening, and accumulation of chlorophyll and anthocyanin (Nagatani *et al*., 1993, Whitelam *et al*., 1993, Neff *et al*., 2000). The control of HY5 by COP1 is one of the primary mechanisms by which a plant decides between skotomorphogenesis and photomorphogenesis (Ang and Deng, 1994, Ang *et al*., 1998, Osterlund *et al*., 2000b, Holm *et al*., 2002, Lau and Deng, 2012).

Many downstream components in light signal transduction have been reported and characterized as involved in photomorphogenic growth and development (Jiao *et al*., 2007, Lee *et al*., 2007). A group of negative regulators known as COP/DET/FUS works downstream of numerous photoreceptors to suppress photomorphogenic development and light-induced gene expression (Wei and Deng, 1999, Jiao *et al*., 2007, Lau and Deng, 2012). COPI acts as a ubiquitin ligase and degrades the photomorphogenesis-promoting factors such as HY5, HYH, LAFl, HFR1 and BBX21 etc. in the dark (Osterlund *et al*., 2000b, Holm *et al*., 2002, Seo *et al*., 2003, Jang *et al*., 2005, Yang *et al*., 2005, Xu *et al*., 2016, Blanco-Touriñán *et al*., 2020). Recent reports have shown that COP1 interacts with SPA1, which facilitates the modulation of the proteasome-mediated degradation of HY5, HFR1, and LAFl (Saijo *et al*., 2003, Seo *et al*., 2003). ELONGATED HYPOCOTYL5 (HY5) is a bZIP transcription factor, a critical regulator of light-mediated seeding photomorphogenic growth (Oyama *et al*., 1997, Chattopadhyay *et al*., 1998b, Gangappa and Botto, 2016, Burko *et al*., 2020). The *hy5* mutant seedlings have a partially etiolated hypocotyl phenotype with reduced chlorophyll and anthocyanin accumulation in white light and different wavelengths of light such as red, far-red, and blue light in addition to having more lateral roots compared with wild-type plants (Koornneef *et al*., 1980, Oyama *et al*., 1997, Chattopadhyay *et al*., 1998b). Moreover, HY5 also functions in UV-B light as a positive regulator of UV-B signalling and (Binkert *et al*., 2014a). HY5 functions cooperatively and antagonistically with various other transcription factors such as HYH, HFR1, CAM7, MYC2, GBF1, TZP and many of the BBX class of proteins either by acting as transcriptional co-activators or co-repressors (Holm *et al*., 2002, Datta *et al*., 2008, Gangappa *et al*., 2013a, Ram and Chattopadhyay, 2013, Abbas *et al*., 2014, Binkert *et al*., 2014b, Gangappa and Botto, 2016, Chakraborty *et al*., 2019, Song *et al*., 2020, Job and Datta, 2021, Li *et al*., 2022).

In Arabidopsis, bHLH transcription factors such as MYC2, MYC3, and MYC4 have been shown to play a vital role in inducing defense responses against herbivory (Dombrecht *et al*., 2007, Chico *et al*., 2014, Zhang *et al*., 2014, Wang *et al*., 2021). MYC2, MYC3, and MYC4 are basic helix-loop-helix transcription factors that interact with Jasmonate Zim-domain proteins and are their direct targets. These TFs have been demonstrated to work together to govern Arabidopsis growth and development (Gao *et al*., 2016). MYC3 is a JAZ-interacting transcription factor that acts with MYC2 and MYC4 to activate JA-responses. Recent studies have also highlighted the importance of MYC2 and MYC4 transcription factors in the regulation of secondary cell wall thickening in response to BL and RL by regulating the expression of genes involved in the secondary cell wall thickening (Zhang *et al*., 2018, Luo *et al*., 2022).

Among MYCs, the role of MYC2 in Arabidopsis seedling development is very well established. MYC2 is a key regulator of seedling photomorphogenesis and light-induced gene expression (Yadav et al., 2005, Gangappa et al., 2010, Sethi et al., 2014, Maurya et al., 2015, Chakraborty et al., 2019, Srivastava et al., 2022, Ojha et al., 2023). While the role of MYC2 in light-mediated seedling development has been studied to some extent, that of MYC3 and MYC4 remains unknown. In this study, we focus on linking the function of light-dependent regulators COP1 and HY5 with MYC2/3/4 in seedling development in Arabidopsis. The genetic and biochemical interaction studies between COP1 and HY5 have greater significance in MYC2/3/4 to control growth in the seedling stage. We also have revealed the stability of MYC3 and MYC4 mediated through COP1 under both light and dark conditions. Our research findings strongly suggest that MYC3 and MYC4 genetically and physically interact with COP1 and HY5 to regulate seedling photomorphogenesis.

## Results

### MYC4 is the negative regulator, while together with MYC3, it promotes seedling photomorphogenesis

To unravel the role of MYC3 and MYC4 in Arabidopsis seedling development, we identified homozygous T-DNA inserted mutant lines for *myc3 (myc3-1)* and *myc4* (*myc4-1)* (Figure S1a, b) and confirmed that they are null alleles as there was no full-length transcript was detected (Figure S1c). We used these alleles for further analysis. Analysis of six-day-old *myc3* mutants grown under SD in white light (WL), blue light (BL), red light (RL), and far-red light (FRL) and also in constant dark (DD) showed no defective hypocotyl length compared to wild-type (Col-0) (Figure 1a-j), suggesting that MYC3 does not control hypocotyl growth independently on its own. However, when we analyzed *myc4* mutant, they showed a significant reduction in hypocotyl length in WL and different wavelengths of light (Figure 1a-h) but not in the DD conditions (Figure 1i, j). Notably, *myc4* mutants showed hypersensitive hypocotyl growth under all the light conditions tested (Figure 1a-h), suggesting that MYC4 likely inhibits photomorphogenesis independent of the wavelength of light.

**Figure 1.**
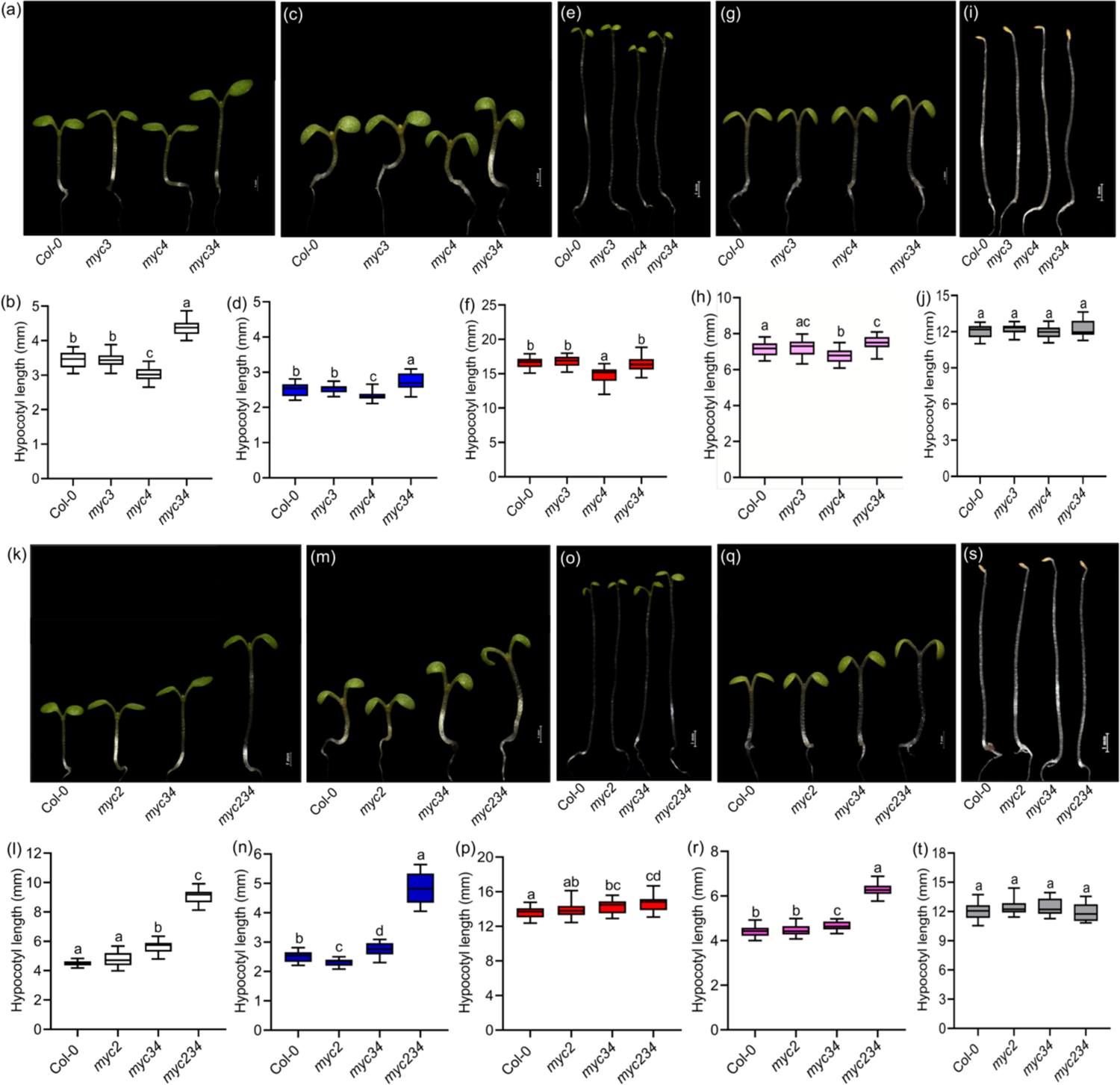
MYC4 inhibits seedling photomorphogenic, while together with MYC3 and MYC2, it strongly enhances photomorphogenic growth. (a-j) Representative seedling images and hypocotyl lengths, respectively, of six-day-old Col-0, *myc3, myc4, and myc34* double mutant seedlings were grown under 22°C SD in WL (a and b), BL (c and d), RL (e and f), FRL (g and h) and DD (i and j). (k-t) Representative seedling images and hypocotyl lengths of six-day-old Col-0, *myc2, myc34,* and *myc234* triple mutant seedlings grown under 22°C SD, respectively, in WL (k and l), BL (m and n), RL (o and p), FRL (q and r) and DD (s and t). Box-whisker plots represent mean±SD. Different letters in the box plot indicate a significant difference (one-way ANOVA with Tukey’s HSD test, P < 0.05, n ≥ 25 seedlings).

MYCs function partially redundantly and additively in controlling defense responses against herbivores and pathogens and in regulating secondary cell-wall thickness (Zhang *et al*., 2018, Van Moerkercke *et al*., 2019, Zhang *et al*., 2020). We wanted to see if MYC3 and MYC4 genetically interact to control seedling hypocotyl growth. We generated a homozygous *myc3myc4* double mutant (hereafter called *myc34*) through a genetic cross and characterized its hypocotyl phenotype to address this. Unexpectedly, *myc34* hypocotyl length was significantly longer than *myc4* and *myc3* and even to Col-0 in WL (Figure 1a, b). Like WL, *myc34* double mutants showed a significant increase in the hypocotyl length than the *myc4* in BL, RL and FRL (Figure 1c-h), suggesting that *myc3* mutation can override the effect of *myc4* mutation. However, in the dark, the *myc34* hypocotyl length was comparable to parental genotypes (Figure 1i, j). Similarly, when we analyzed *myc2myc3* (*myc23*) and *myc2myc4* (*myc24*) double mutant hypocotyl lengths in various lights, it was found that *myc23* and *myc24* double mutants had hypocotyl lengths taller than Col-0 as seen in WL, and similar to Col-0 in BL, RL and FR (Figure S2a-h) but not in DD (Figure S2i, j). These results imply that even though MYC2 and MYC4 alone function as negative regulators, together with MYC3, they likely act as positive regulators of light-mediated inhibition of seedling hypocotyl growth.

Light differentially controls hypocotyl and cotyledon growth. As MYC3/MYC4 together promote hypocotyl growth inhibition, we wanted to see if they also control cotyledon growth. We examined the cotyledon phenotype of six-day-old Col-0, *myc3*, *myc4* and *myc34* seedlings grown in SD photoperiods. Cotyledon area measurement data from six-day-old seedlings reveal that *myc3* had significantly bigger cotyledons than Col-0 in WL and FR, while it was comparable to Col-0 in BL and RL (Figure S3a-e). The cotyledon area of the *myc4* mutant was comparable to Col-0 in WL and across different wavelengths of light (Figure S3a-e). Interestingly, the *myc34* mutants had significantly bigger cotyledons than single mutants (Figure S3a-e). These results suggest that MYC3/MYC4 functions partially redundantly to inhibit cotyledon expansion and differentially regulate photomorphogenic growth in the cotyledon and hypocotyl.

### MYC3/MYC4 genetically interact with MYC2 and together promote seedling photomorphogenesis

Knowing that MYC3 and MYC4 function partially redundantly with MYC2 in controlling defense responses and other aspects of the growth and development (Zhang *et al*., 2018, Van Moerkercke *et al*., 2019, Zhang *et al*., 2020), we were intrigued to know the genetic interaction among all three MYCs in seedling photomorphogenic growth. We identified a *myc2myc3myc4* homozygous triple mutant (hereafter called *myc234*) by crossing the *myc34* double mutant with the *myc2*, *jin1-2* allele (Lorenzo *et al*., 2004). Measuring the hypocotyl phenotype of six-day-old seedlings grown in WL under SD photoperiod suggests that, surprisingly, the *myc2/jin1-2* mutation further enhanced the hypocotyl length of the *myc34* double mutant (Figure 1k, l). At the same time, *myc34* showed significantly longer hypocotyls than Col-0, as observed above, while *myc2* mutant hypocotyl length was comparable to Col-0 under SD (Figure 1k, l). Like WL, the hypocotyl length of the *myc234* triple mutant under BL showed significantly elongated hypocotyl than the *myc34* double mutant (Figure 1m, n). However, as reported, *myc2* showed significantly shorter hypocotyl than Col-0 in BL (Figure 1m, n). In RL, the triple mutant hypocotyl length was comparable to Col-0 (Figure 1o, p), while in FRL, it showed significantly longer hypocotyls than the double mutants and Col-0 (Figure 1q, r). In DD, the *myc234* triple mutant hypocotyl length was similar to parental genotypes (Figure 1s, t). These results reveal that MYC3/MYC4 interact with MYC2 and exhibit unequal genetic redundancy to inhibit light-mediated seedling hypocotyl growth in WL, BL and FRL.

### MYC2/MYC3/MYC4 together regulate anthocyanin and chlorophyll accumulation

Besides inhibiting hypocotyl growth, seedling photomorphogenesis also accompanies an increased accumulation of photopigments such as anthocyanin and chlorophyll. As *myc234* triple mutants showed a strong light-insensitive response to hypocotyl growth inhibition, we wanted to know if the accumulation of photopigments is also altered. Analysis of anthocyanin content from six-day-old seedlings revealed that *myc2* and *myc4* accumulated slightly but significantly more anthocyanin in WL, BL and RL (Figure S4a-c); however, the *myc3* mutant accumulated slightly more anthocyanin in RL but not in other light conditions (Figure S4c). The anthocyanin content in the *myc34* double mutants is largely comparable to Col-0 (Figure S4a-d). Interestingly, consistent with the reduced light sensitivity of hypocotyl, in the *myc234* triple mutant, anthocyanin content was significantly reduced than Col-0 in WL and different wavelengths of light (Figure S4a-d), suggesting that MYC2/MYC3/MYC4 together act to promote anthocyanin accumulation.

Unlike anthocyanin, the triple mutants had significantly less chlorophyll content in WL and BL (Figure S4e, f) but not in RL and FRL (Figure S4g, h). The *myc34* had chlorophyll that was largely similar to Col-0 in WL and other wavelengths of light (Figure S4e-h). However, the *myc2* and *myc4* mutants accumulated significantly more chlorophyll than Col-0 in WL and BL (Figure S4e, f). but not in RL and FRL (Figure 4Sg, h). These data suggest that MYC2/MYC3/MYC4 differentially regulate chlorophyll and anthocyanin accumulation in a wavelength-dependent manner.

### MYC2/MYC3/MYC4 modulates the expression light-inducible genes

MYC2 has been shown to inhibit the expression of light-inducible genes such as *CAB1, RBCS-1A,* and *CHS* involved in photosynthetic reaction and anthocyanin biosynthesis (Yadav *et al*., 2005). We are curious to know how *MYCs* individually and together control the expression of light-inducible genes. The expression of these genes is significantly reduced in the *myc234* triple mutant than Col-0 when we exposed constant dark-grown seedlings to light for 4h (Figure 2a-f). Consistent with reduced anthocyanin and chlorophyll accumulation, the *myc234* triple mutants showed compromised expression of various light-inducible genes such as *CHLOROPHYLL A/B BINDING 1* (*CAB1*), *RIBULOSE BISPHOSPHATE CARBOXYLASE SMALL SUBUNIT-1A (RBCS-1A*) and *EARLY LIGHT-INDUCIBLE PROTEIN 2* (*ELIP2*), involved in the photosynthetic reaction; and *CHALCONE SYNTHASE* (*CHS*) and *CHALCONE ISOMERASE* (*CHI*), which code for enzymes involved in anthocyanin biosynthesis (Figure 2a-e). Moreover, the key transcription factor *HY5*, which also directly binds and activates *CAB1*, *RBCS-1A*, *ELIP2*, *CHS* and *CHI,* showed significant regulation in the *myc234* triple mutant than Col-0 (Figure 2f). In the *myc34* double mutants, the expression of these genes is largely comparable to Col-0 (Figure 2a-e). Interestingly, the expression of *CAB1*, *RBCA-1A*, *ELIP2*, *CHS* and *CHI* was significantly elevated in the *myc2* mutant than Col-0 (Figure 2a-e), while in *myc4* it was largely similar to Col-0 in 4h light treated seedlings. Notably, some genes, such as *ELIP2*, *CHS, and CHI,* showed significant upregulation in the *myc3 and myc4* single mutants compared to Col-0 in the dark (Figure 2c-e). These gene expression data suggest that MYCs differentially regulate the expression of light-inducible genes.

**Figure 2.**
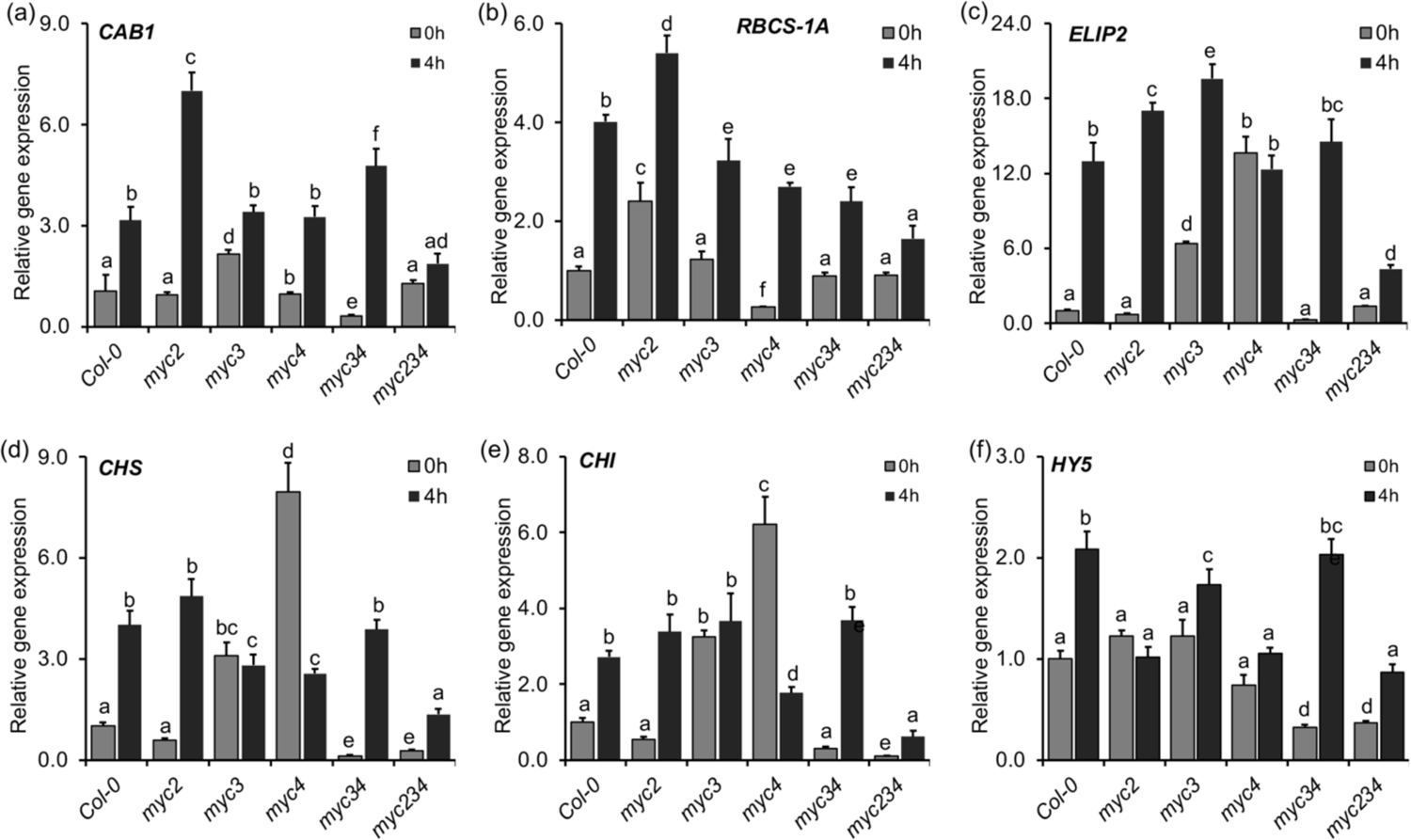
Expression of light-inducible genes in various *myc* mutants. Expression of light-inducible genes *CAB1* (a)*, RBCS-1A* (b)*, ELIP2* (c)*, CHS* (d)*, CHI* (d) and *HY5* (f) in six-days-old seedlings of Col-0*, myc2, myc3, myc4, myc34, and myc234* grown in constant dark for six-days and 4-h light (WL) treated seedlings. The bars represent the mean±SD (n= three biological replicates). The data were first normalized with *EF1α* and then calculated fold-change against wild-type 0h as 1. Different letters above the bars indicate a significant difference from other genotypes or treatments (one-way ANOVA with Tukey’s HSD test, P < 0.05, n ≥ 25 seedlings).

### MYC4 protein stability is regulated by light

As MYC3 and MYC4 regulate seedling photomorphogenic growth, we wanted to know if light regulates their gene expression and protein accumulation. We carried out our qRT-PCR analysis from six-day-old seedlings and found that both *MYC3* and *MYC4* gene expression is significantly upregulated in WL, BL RL and FR compared to constant dark-grown seedlings (Figure S5a). Also, when we analyzed their expression in different tissues, we found that *MYC3* and *MYC4* ubiquitously expressed across different tissue types (Figure S5b).

Consistent with the increased accumulation of *MYC4* transcript in light, our immunoblot assays using *35S:MYC4-GFP* (Fernandez-Calvo *et al*., 2011) reveal that MYC4 is more stable in the light as the protein stability was higher in WL-grown SD, LD and LL photoperiods than DD (Figure 3a). Compared to SD, LD-grown seedlings accumulated slightly more MYC4 protein, while constant light (LL) grown seedlings accumulated a similar protein level to SD (Figure 3a). Similarly, when DD-grown seedlings were shifted to WL for different durations, light-treated seedlings accumulated ∼two-fold more MYC4 protein than the DD-grown seedlings (Figure 3b). Moreover, compared to ZT0, MYC4 protein stability was higher during the daytime (ZT2-ZT8) and started declining at the end of the day (ZT12-ZT16) (Figure 3c). Besides, when we checked the MYC4 protein stability under different light conditions, its stability was higher in WL and BL compared to RL, FRL and DD (Figure 3d). Together, these immunoblot results reveal that light promotes the stability of MYC4 protein accumulation. Further, we checked the stability of MYC3 using *35S:MYC3-GFP* transgenic line (Fernandez-Calvo *et al*., 2011). Unlike MYC4, we could not detect MYC3 protein under any tested conditions, including different wavelengths and photoperiods (Figure S6a-d), suggesting that MYC3 protein is probably highly unstable, indicating MYC3 function could be under tight control. Its activity is likely being controlled at the posttranslational level by E3 ubiquitin ligase(s), this could be one of the factors contributing to the complex genetic interactions observed among MYCs.

**Figure 3.**
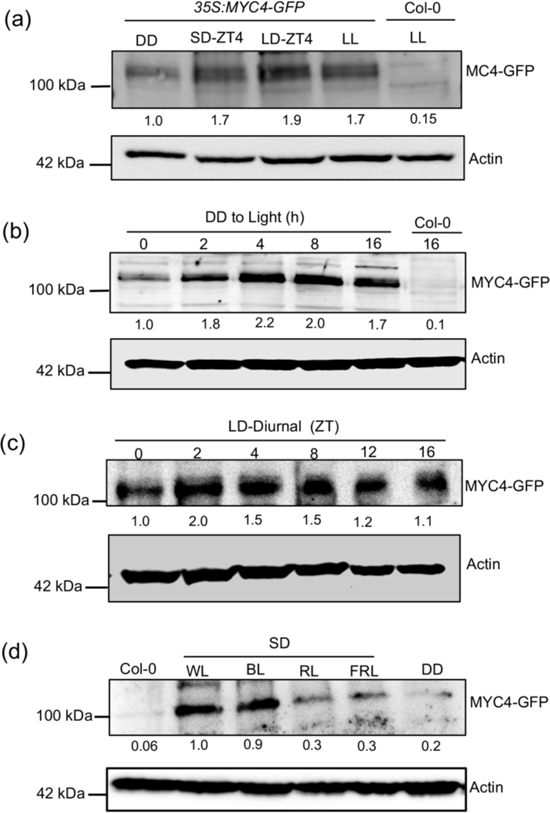
MYC4 protein stability is regulated by light (a) Immunoblot analysis from six-day-old seedlings of *35S:MYC4-GFP* transgenic line in constant dark (DD), SD (ZT4), LD (ZT4) and constant light (LL). (b) Immunoblot analysis of *35S:MYC4-GFP* transgenic line. Six-day-old DD-grown seedlings were treated with WL for different time duration as indicated. (c) Immunoblot analysis of six-day-old seedlings of *35S:MYC4-GFP* line grown in WL under LD photoperiods. (d) Immunoblot analysis of *35S:MYC4-GFP* seedlings grown under different monochromatic lights BL, RL and FRL, including WL and DD. In panels (a), (b) and (d), Col-0 seedlings were used as a negative control. The total protein was extracted and then subjected to immunoblot analysis using an anti-GFP antibody. The MYC4-GFP is shown as indicated. The lower panels show the immunoblot of anti-Actin as loading controls. Values underneath the blots are relative protein levels, calculated after normalizing to actin using ImageJ software.

### MYC3 and MYC4 regulate petiole and rosette development

To further know if MYC3 and MYC4 influence growth beyond seedling development, we examined their petiole and rosette phenotypes. The *myc3* mutant had a similar rosette size, while *myc4* had significantly smaller rosettes than Col-0 under SD and LD photoperiods (Figure S7a-c). Interestingly, the *myc34* double mutants had slightly but significantly increased rosette diameter under SD and LD (Figure S4a-c). In line with the increased rosette diameter, the *myc3* mutant had petiole length similar to Col-0, while *myc4* had significantly shorter petioles (Figure S4c, d). However, the petiole length of *myc34* double mutants was significantly increased (Figure S4c, d). These results suggest that similar to regulating hypocotyl growth, MYC3/MYC4 probably show unequal redundancy in regulating rosette size and petiole growth.

### MYC3 and MYC4 genetically interact with HY5 and suppress its function

HY5, a potent activator of seedling photomorphogenesis, functions downstream to all the known photoreceptors (Oyama *et al*., 1997, Chattopadhyay *et al*., 1998b). We were interested to see if MYC3 and MYC4 genetically interact with HY5 in regulating seedling hypocotyl growth. We generated *myc3hy5* and *myc4hy5* double mutants and investigated their phenotypes. Measurement of hypocotyl lengths revealed that *hy5* showed long and partially etiolated hypocotyls in WL compared to Col-0 as reported (Ang and Deng, 1994, Chattopadhyay *et al*., 1998b) (Figure 4a, b). However, *myc3* and *myc4* mutations significantly suppressed the *hy5* hypocotyl phenotype as the hypocotyl length of *myc3hy5* and *myc4hy5* was significantly reduced than *hy5* (Figure 4a, b). Similarly*, myc3hy5* and *myc4hy5* double mutants displayed significantly shorter hypocotyls than *hy5* in BL, RL, and FRL conditions (Figure 4c-h), suggesting that MYC3 and MYC4 genetically interact with HY5 and probably function antagonistically to control hypocotyl growth. The suppression of *hy5* mutant hypocotyl length by *myc3* and *myc4* is also consistent with the genetic interaction observed for MYC2 and HY5 (Chakraborty et al., 2019), wherein *myc2* could strongly suppress the long hypocotyl phenotype of *hy5*. Consistent with the unequal genetic redundancy observed in *myc34* double mutants, we wanted to test the effect of *myc34* mutations together in modulating *hy5* mutant hypocotyl growth. We generated a *myc3myc4hy5* (*myc34hy5*) triple mutant and measured hypocotyl length in WL and different monochromatic lights. Our data revealed that the hypocotyl length of the *myc34hy5* triple mutant was slightly but significantly longer than the *myc3hy5* and *myc4hy5* double mutants (Figure 4a-h), suggesting that when both MYC3 and MYC4 are absent, their ability to suppress *hy5* hypocotyl phenotype is reduced than when any one of them is absent.

**Figure 4.**
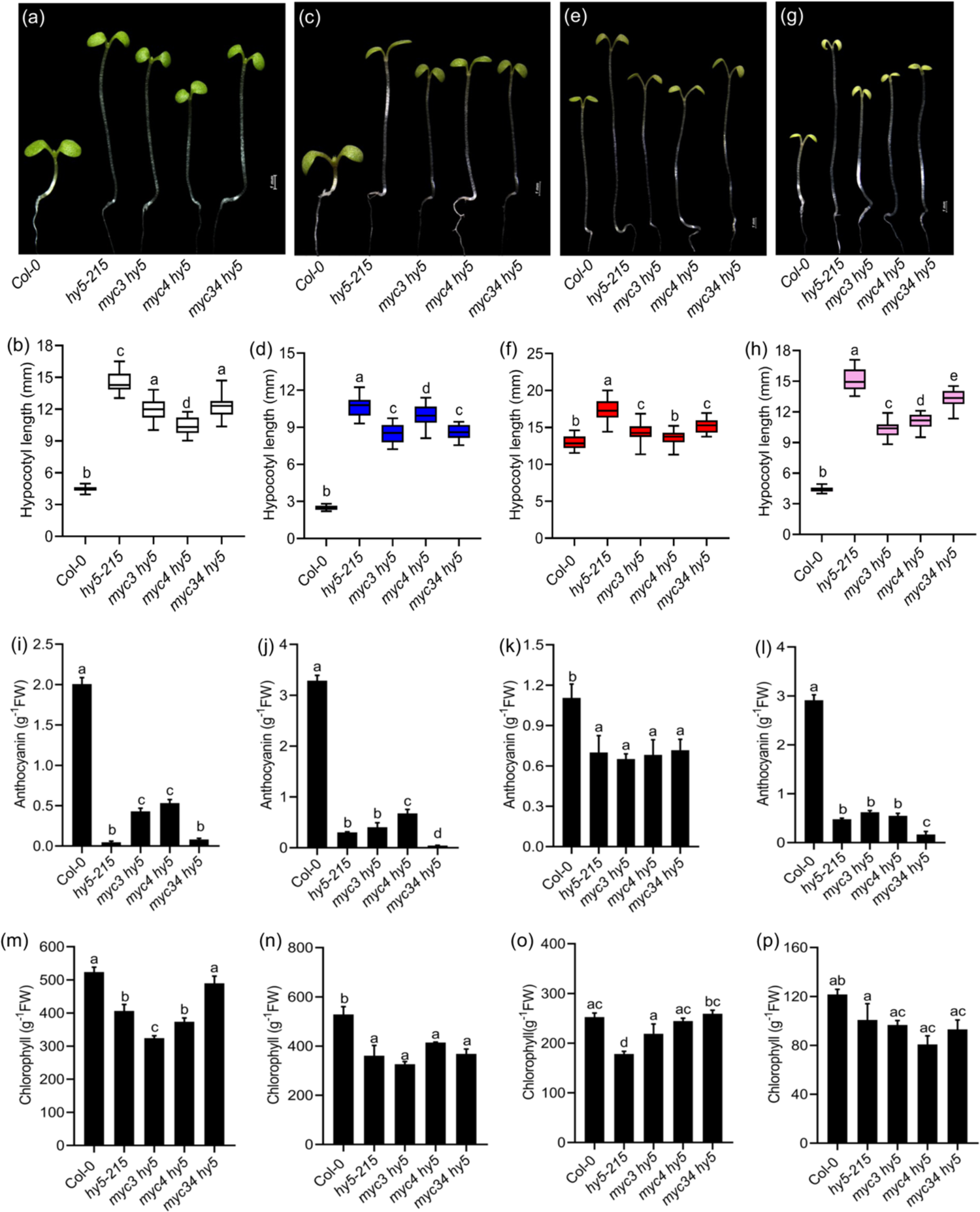
MYC3/MYC4 genetically interact with HY5 to regulate seedling photomorphogenic growth. (a-h) Representative seedling pictures and the hypocotyl length of six-day-old Col-0, *hy5-215, myc3hy5, myc4hy5, myc34 hy5* triple mutant seedlings grown in WL (a and b), BL (c and d), RL (e and f), FRL (g and h) under SD photoperiod. (i-l) Quantification of anthocyanin content in the indicated genotypes shown grown in SD photoperiod for six days in WL (i), BL (j), RL (k), and FRL (i). (m-p) Chlorophyll content of six-day-old seedlings of mentioned genotypes grown in WL (m), BL (n), RL (o) and FRL (p) conditions. The data shown is mean± SD, ≥20 seedlings). Different letters above the bar chart indicate a significant difference (one-way ANOVA with Tukey’s HSD test, P < 0.05).

Similar to the effect seen in controlling hypocotyl growth, *myc3* and *myc4* mutations also significantly suppressed low anthocyanin levels of *hy5* in WL and BL (Figure 4i, j). However, in the *myc34hy5* triple mutant, anthocyanin levels were comparable to *hy5* (Figure 4i, j). In RL, anthocyanin levels in the *myc3hy5*, *myc4hy5* and *myc34hy5* were largely similar to the *hy5* mutant (Figure 4k). In FRL, the anthocyanin levels in the *myc3hy5* and *myc4hy5* were comparable to *hy5*, while it is further reduced than *hy5* in the *myc34hy5* triple mutants (Figure 4l). Similarly, *myc3hy5*, *myc4hy5,* and *myc34hy5* accumulated chlorophyll content similar to *hy5* in all the light conditions, including WL (Figure 4m-p). However, *myc34* suppressed the low chlorophyll content of *hy5* in WL and RL but not in BL and FRL (Figure 4m-p), further suggesting that MYC3 and MYC4 probably influence HY5 function differentially in a light-dependent manner to regulate anthocyanin and chlorophyll accumulation.

### MYCs act as rate-limiting factors for the optimal accumulation of HY5 protein

As MYCs modulate HY5-mediated hypocotyl growth, we were curious to know if this is through regulating HY5 protein levels. We used HY5-specific native antibodies and carried out immunoblotting assays (Figure S8). Six-day-old seedlings grown in WL under SD revealed that in *myc2*, *myc3, myc4* single mutant, and *myc34* double mutants, HY5 protein stability is comparable to Col-0 (Figure 5a). However, in *myc234*, its stability is reduced by at least two folds (Figure 5a). Like SD, seedlings grown in WL under LD also showed reduced HY5 stability in *myc234* triple mutants (Figure 5b). but not in the single and double mutants of *MYCs*, suggesting that MYC2/MYC3/MYC4 play a key role in maintaining optimal HY5 protein levels. In BL-grown seedlings, we saw a moderate to drastic reduction in the stability of HY5 protein in *myc34* and *myc234* mutants (Figure 5c). However, the HY5 protein stability was slightly reduced in the *myc2* mutant but was comparable to Col-0 in *myc3* and *myc4* single mutants. Contrarily, in the RL and FRL conditions, the HY5 protein stability was ∼ two-fold elevated in the *myc34* double and *myc234* triple mutants compared to Col-0 and the single mutants (Figure 5d, e). Notably, HY5 protein stability was slightly elevated in the *myc2*, *myc3* and *myc4* single mutants, compared to Col-0 (Figure 5d, e). These data suggest that MYC2/MYC3/MYC4 differentially regulate HY5 protein stability in a light wavelength-dependent manner.

**Figure 5.**
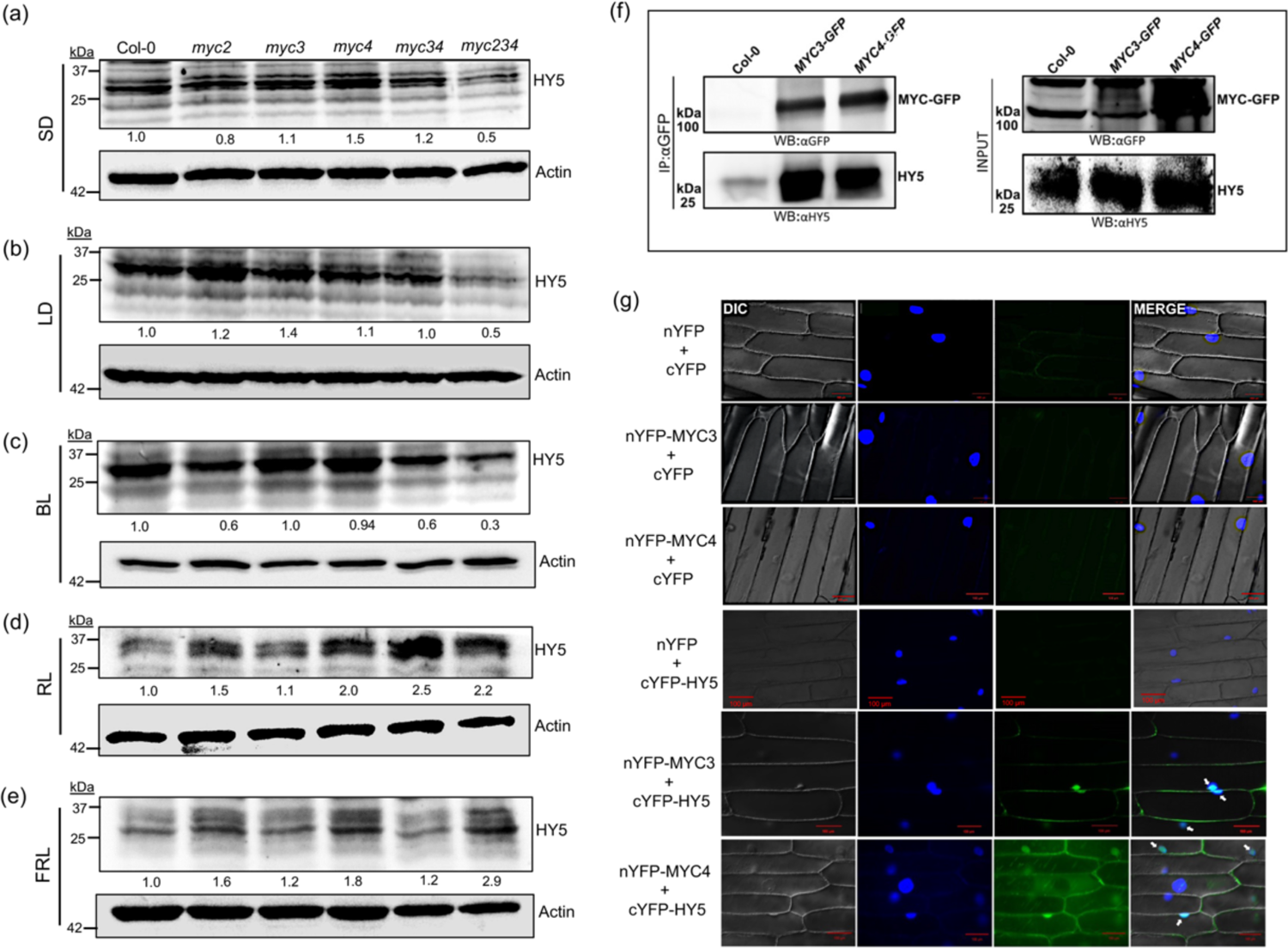
MYC2/MYC3/MYC4 regulate HY5 protein stability, probably through physical interaction (a-e) Immunoblot analysis for checking HY5 protein levels using native HY5 protein antibody in six-day-old seedlings of Col-0, *myc2, myc3, myc4, myc34* and *myc234* triple mutant grown in WL under SD (a) and LD (b); and under SD in BL (c), RL (d) and FRL (e). Actin blots were shown for loading control and were also used for normalization to calculate the relative protein levels shown underneath each immunoblot. The experiments were repeated twice, and similar results were obtained. (f) Immunoblot showing coimmunoprecipitation of HY5 when MYC3-GFP or MYC4-GFP proteins were immunoprecipitated using an anti-GFP antibody. The *35S:MYC3-GFP 35S:MYC4-GFP* transgenic lines grown in WL under LD for six days were used. As MYC3 is highly unstable, *35S:MYC3-GFP* seedlings treated with MG132-treated were used for the assay. The immunoprecipitated complex was resolved by SDS-PAGE. Both the input and IP were probed with antibodies to HY5. The tissue harvested at ZT4 was used for Co-IP assays. Wild type (Col-0) was used as a negative control. (g) BiFC assay suggests physical interaction between MYC3/MYC4 and HY5. All the constructs containing nYFP and cYFP were co-transformed into onion epidermal cells. nYFP-MYC3/cYFP-HY5 and nYFP-MYC4/cYFP-HY5 combination show positive signals for interactions. The left panel image shows the bright field image (DIC), the middle panel shows the nucleus staining by DAPI, and the next panel shows the YFP channel produced by the reconstruction of YFP. The right panel image shows the merged image. The Arrows indicate the reconstituted YFP signal in the nuclei.

### MYC3/MYC4 physically interact with HY5

As MYC3 and MYC4 genetically interact and regulate HY5 protein stability to control seedling photomorphogenic growth, we wanted to check if MYC3/MYC4 physically interact with HY5 and regulate its protein function and protein stability. To address this, we performed an In vivo co-immunoprecipitation assay using six-day-old *35S:MYC3-GFP* seedlings treated MG132 and *35S:MYC4-GFP* transgenic lines grown in WL under LD. Our immunoblot data show that when either MYC3-GFP or MYC4-GFP was immunoprecipitated using an anti-GFP antibody, we could detect HY5 protein being pulled down in immunoprecipitated complex along with MYC3-GFP and MYC4-GFP, as detected using an anti-HY5 antibody (Figure 5f). However, when we used Col-0 as a negative control, HY5 protein was not detectable in the immunoprecipitated complex (Figure 5f), suggesting that HY5 could physically associate with MYC3 and MYC4. These results were further confirmed by BiFC assay in the onion epidermal cells. When we co-infiltrated cYFP-HY5 with either nYFP-MYC3 or nYFP-MYC4, we could detect a YFP fluorescence in the nucleus, as revealed by microscopy (Figure 5g). However, we could not detect fluorescence when cYFP-HY5/nYFP, cYFP/nYFP-MYC3, cYFP/nYFP-MYC4 or nYFP/cYFP combinations were co-infiltrated (Figure 5g).

### MYC3 and MYC4 genetically interact with COP1 and modulate its function

COP1 is a critical repressor of the photomorphogenesis (Deng *et al*., 1991). It ubiquitinates and degrades many positive regulators of seedling photomorphogenesis, such as HY5, HYH, LAF1, HFR1, BBX proteins, etc. (Osterlund *et al*., 2000a, Holm *et al*., 2002, Seo *et al*., 2004, Lau and Deng, 2012). As MYC3 and MYC4 interfere with the HY5 function, we wanted to know if they have any functional connection with COP1. We generated *myc3cop1-4*, *myc4cop1-4* double mutants and *myc34cop1-4* triple mutants to address. Measurement of hypocotyl length from six-day-old constant dark-grown (DD) seedlings revealed that *myc3cop1-4* had hypocotyl length largely similar to the *cop1-4* single mutant, while *myc4cop1-4* had subtle but significantly shorter hypocotyls than *cop1-4* (Figure 6a, b). Similarly, the *myc34cop1-4* triple mutant hypocotyl length was significantly shorter than *cop1-4* but comparable to the *myc4cop1-4* double mutant (Figure 6a, b). Like DD, six-day-old WL-grown *myc3cop1-4 and myc4cop1-4* seedlings under SD photoperiod had significantly shorter hypocotyls than *cop1-4,* while the *myc34cop1-4* triple mutant hypocotyl length was comparable to the *myc3cop1-4* double mutant (Figure 6c, d). In BL, *myc3cop1-4* hypocotyl length was comparable to *cop1-4,* while the *myc4cop1-4* and *myc34cop1-4* had significantly shorter hypocotyls than *cop1-4* (Figure 6e, f). In RL, the hypocotyl length of *myc3cop1-4*, *myc4cop1-4* and *myc34cop1*-4 triple mutants was comparable to *cop1-4* (Figure g, h). In. FRL, the *myc3 cop1-4* had significantly shorter hypocotyls than *cop1-4,* while *myc4cop1-4* and *myc34cop1-4* hypocotyl length was comparable to *cop1-4* (Figure 6i, j). These results indicated that MYC3 and MYC4 likely enhance COP1 function to promote hypocotyl elongation in dark and light.

**Figure 6.**
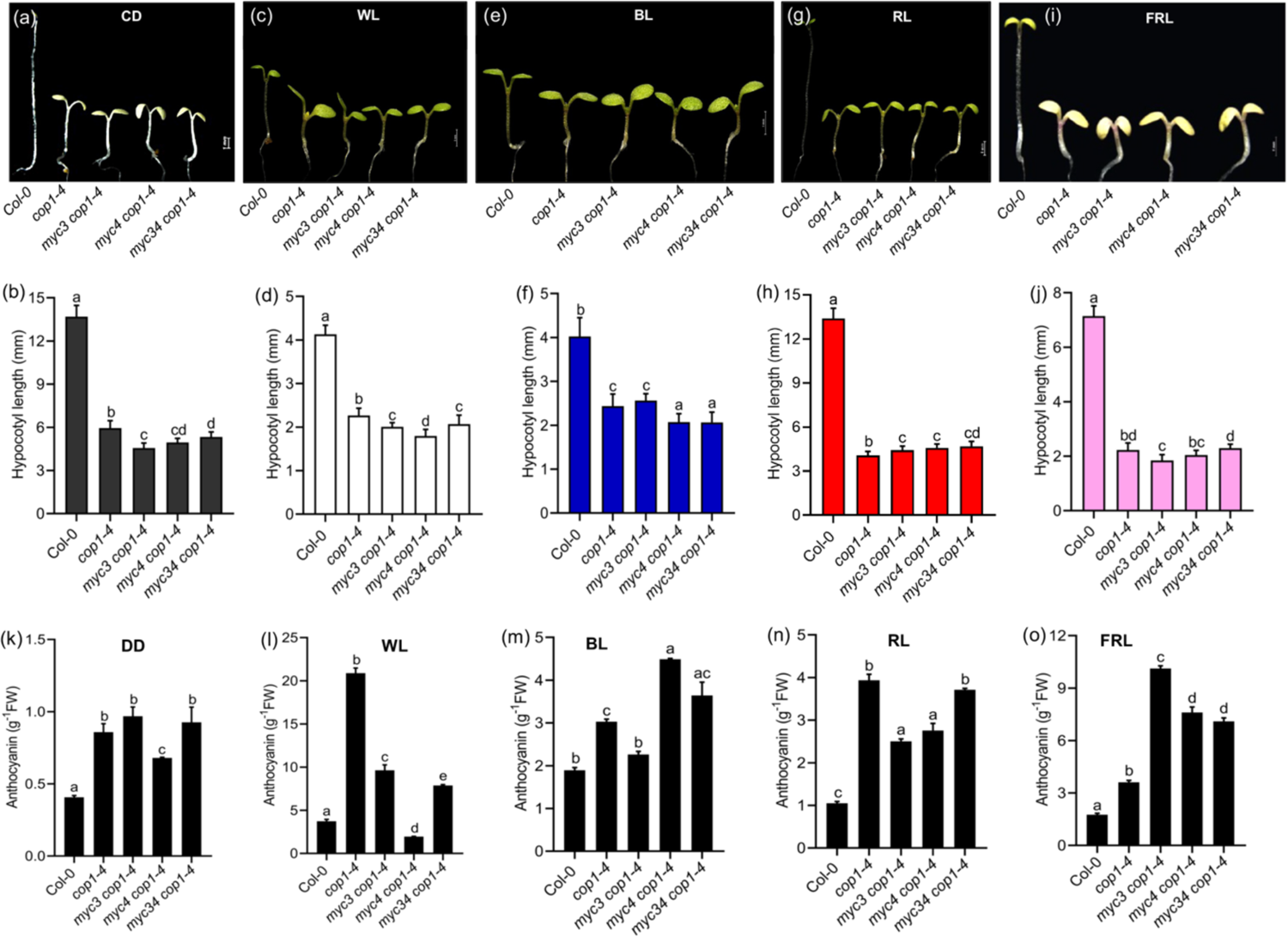
Genetic interaction between MYC3/MYC4 and COP1 to control regulation of photomorphogenesis. (a-j) Representative seedling images and measured hypocotyl length of Col-0, *cop1-4*, *myc3cop1-4, myc4cop1-4, myc34cop1-4* triple mutant seedlings grown for six days in constant dark (a and b); and under SD photoperiod in WL (c and d), BL (e and f), RL (g and h) and FRL (i and j). (k-o) Anthocyanin contents in the indicated genotypes grown in SD photoperiod for six days in constant dark (k) and under SD in WL (l), BL (m), RL (n), and FRL (o). The data shown is mean± SD, ≥20 seedlings). Different letters above the bar chart indicate a significant difference (one-way ANOVA with Tukey’s HSD test, P < 0.05).

### MYC3 and MYC4 regulate COP1-mediated anthocyanin accumulation in dark and light

The various alleles of *cop1* mutant accumulate elevated anthocyanin in addition to suppressing hypocotyl growth. To know if MYC3 and MYC4 interfere with COP1 in controlling anthocyanin accumulation, we quantified anthocyanin content in DD, WL, and different nonchromatic light-grown seedlings. As expected, the anthocyanin content in *cop1-4* was significantly higher than that of Col-0 in DD. The *myc3cop1-4* and *myc34cop1-4* mutants were comparable to *cop1-4* (Figure 6k). However, in the *myc4cop1-4*, anthocyanin levels are slightly lower than *cop1-4* but significantly more than Col-0 (Figure 6k). In WL-SD-grown seedlings, *cop1-4* accumulated significantly more anthocyanin than Col-0 (Figure 6l). However, in the *myc3cop1-4, myc4cop1-4* and triple mutants, anthocyanin content was significantly reduced than *cop1-4* (Figure 6l). However, in BL, the anthocyanin content of *myc4cop1-4* and *myc34cop1-4* was significantly elevated compared to that of *cop1-4*, while it was similar to Col-0 (Figure 6m). In the RL, anthocyanin content in *myc3cop1-4, myc4cop1-4* and *myc34cop1-4* mutants was significantly elevated than the *cop1-4*. However, in the triple mutant, there was more than in both the single mutants (Figure 6n). Like BL, the anthocyanin content *cop1-4* was further enhanced in double and triple mutants under FRL (Figure 6o).

While COP1 is a negative regulator of anthocyanin in the dark and across different wavelengths of light, it differentially regulates chlorophyll accumulation in a light and allele-specific manner. In WL and FRL, *cop1-4* accumulates less chlorophyll than Col-0, while the *myc3cop1-3, myc4cop1-4 cop1-4* double and *myc34cop1-4* triple mutants had similar chlorophyll levels to *cop1-4* (Figure S9a, d). In BL, *cop1-4* accumulated more chlorophyll than Col-0, while *myc3myc4* further enhanced the chlorophyll content of the *cop1-4* mutant (Figure S9b). Like BL, *cop1-4* accumulates more chlorophyll in RL than Col-0 (Figure S9c). However, in the *myc3cop1-3, myc4cop1-4* double and *myc34cop1-4* triple mutant, it was comparable to *cop1-4* (Figure S9b), suggesting that MYC3 and MYC4 interaction with COP1 for the regulation of anthocyanin and chlorophyll is dependent on wavelength of light.

### MYC4 interfere with COP1-mediated degradation of HY5 protein in light

COP1-mediated skotomorphogenic growth depends on the degradation of photomorphogenesis-promoting factors such as HY5, HYH, LAF1, HFR1, BBX proteins, etc. We tested if MYC3/MYC4 regulate HY5 protein stability through COP1. To address this, we grew Col-0, *cop1-4*, *myc3*, *myc4*, *myc3cop1-4, myc4cop1-4* and *myc34cop1-4* genotypes in DD for six days and checked for HY5 protein stability using anti-HY5 antibody. Our data suggest that HY5 protein was elevated in the *cop1-4* mutant background. While in the *myc3cop1-4* mutant, HY5 protein stability was comparable to *cop1-4*, it was further enhanced in the *myc4cop1-4* and *myc34cop1-4* mutants (Figure 7a). When DD-grown seedlings were transferred to WL for 8h, HY5 protein stability was slightly elevated in the *myc34cop1-4* triple mutants, while in *myc3cop1-4* and *myc4cop1-4* double mutants, its stability was comparable to *cop1-4* (Figure 7b). In the *myc3* and *myc4* single mutants, HY5 stability was largely comparable to Col-0 (Figure 7b). In WL-grown seedlings under SD, the HY5 stability was further enhanced in the *myc4cop1-4* and *myc34cop1-4*, but in the *myc3 cop1-4* mutant, it was comparable to *cop1-4* and Col-0 (Figure 7c). Also, in *myc3* and *myc4* mutants, HY5 stability was similar to Col-0 (Figure 7c). These results suggest that MYC4, not MYC3, might help COP1 promote HY5 degradation.

**Figure 7.**
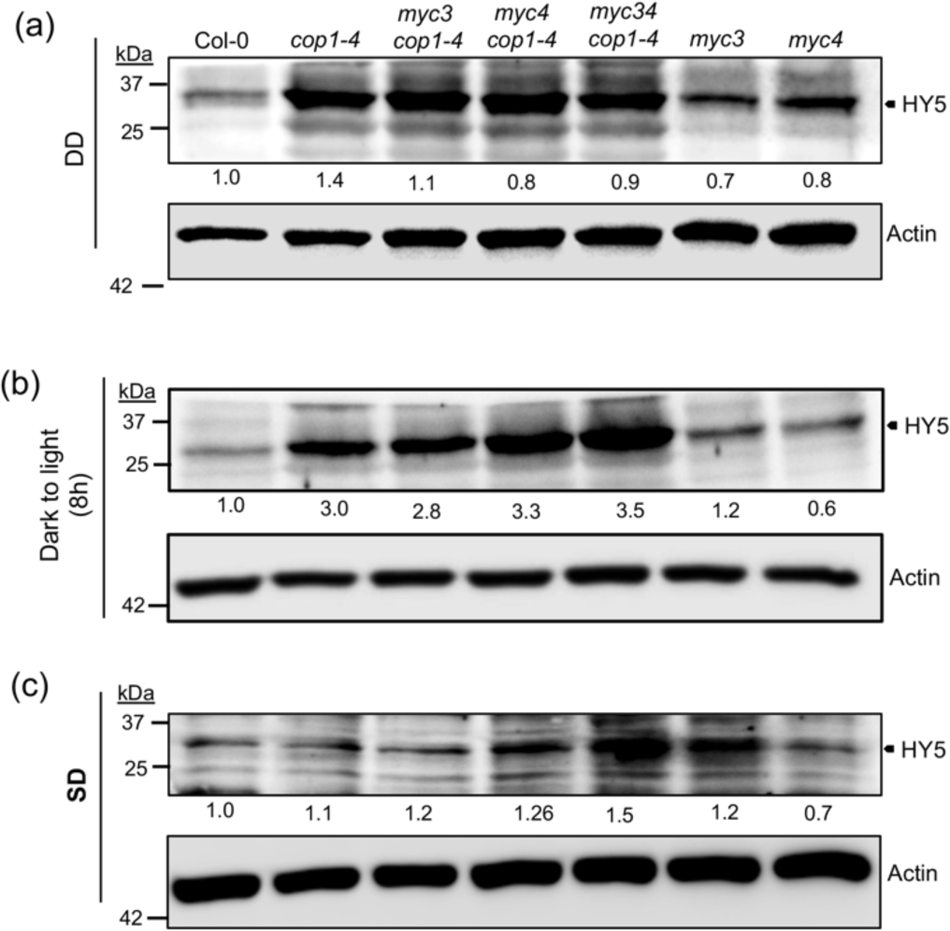
MYC4 probably helps COP1 in the degradation of HY5 and modulation of seedling photomorphogenesis. (a-c) Immunoblot analysis showing HY5 protein stability using native HY5 antibody in Col-0, *cop1-4, myc3cop1-4, myc4cop1-4, myc3myc4cop1-4, myc3* and *myc4* mutants under constant dark (DD)(a), from constant dark to light treated for 8 h (b) and at ZT4 under SD (c). Actin was used as a control for all the blots and normalization. Relative protein levels shown underneath each immunoblot were calculated using ImageJ software. The experiments were repeated two times, and similar results were obtained.

### MYC3 and MYC4 are targeted by COP1 for degradation thorough 26S proteasomal pathway

Our results suggest that MYC3 and MYC4 interfere with COP1-mediated hypocotyl and photo-pigment accumulation. We sought to examine if COP1 regulates MYC3 and MYC4 protein stability. To address this, we introgressed *35S:MYC3-GFP* and *35S:MYC4-GFP* transgenes into the *cop1-4* mutant background and identified homozygous *cop1-4 35S:MYC3-GFP* and *cop1-4 35S:MYC3-GFP* lines. Immunoblot analysis of MYC3 and MYC4 protein stability from six-day-old DD-grown seedlings revealed that MYC3 protein was highly unstable (Figure 8a), while MYC4 protein was moderately stable (Figure 8a), as observed above. Interestingly, in the *cop1-4* mutant background, MYC3-GFP was stabilized (Figure 8a), suggesting that MYC3 is probably under the tight control of COP1. Similarly, MYC4-GFP protein stability was also elevated in the *cop1-4* mutant background compared to the wild-type (Figure 8a), suggesting that COP1 degrades MYC4 protein as well in the dark. Similar to DD, in the *cop1-4* mutant background, MYC3-GFP was stabilized in WL and different wavelengths of light (Figure 8b-e)), suggesting that MYC3 is probably degraded by COP1 in both dark, WL and different monochromatic lights. On the other hand, unlike MYC3, the stability of MYC4-GFP protein in the *cop1-4* mutant was compared to the Col-0 in WL and BL conditions (Figure 8b, c). However, in the RL and FRL conditions, MYC4-GFP protein stability was higher in the *cop1-4* background than Col-0 (Figure 8d, e), suggesting that MYC4 protein degradation by COP1 was wavelength-dependent.

**Figure 8.**
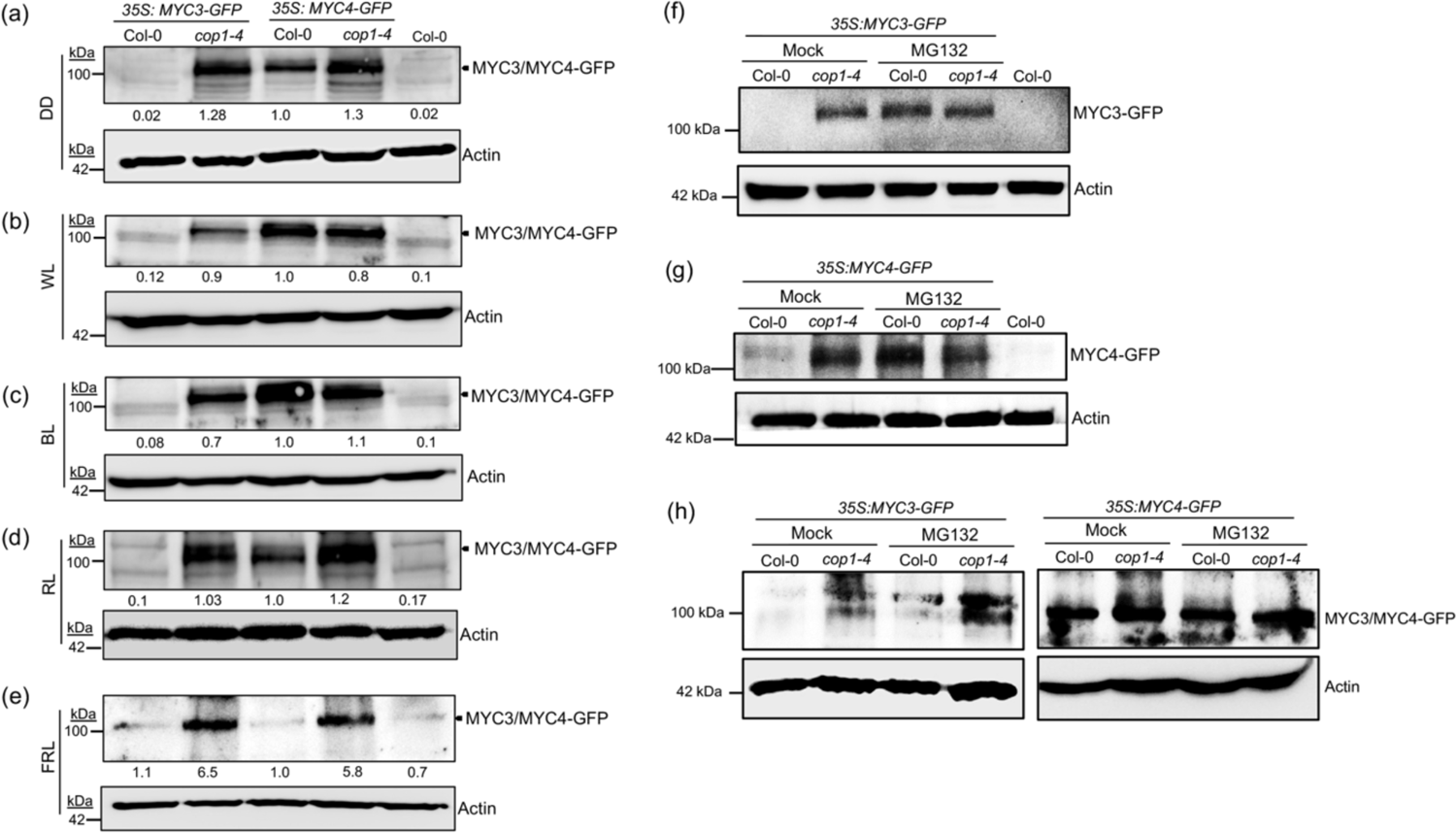
COP1 targets MYC3 and MYC4 for proteasomal degradation (a-e) MYC3 and MYC4 proteins are stabilized by 26S proteasome inhibitor (MG132). Immunoblot analysis showing MYC3/MYC4 protein stability in *35S:MYC3-GFP, cop1-4 35S:MYC3-GFP, 35S:MYC4-GFP* and *cop1-4 35S:MYC4-GFP* seedlings grown under DD (a), and in WL (b), BL (c), RL (d), and FRL (f), under SD photoperiod. (f, g) Immunoblot analysis of MYC3 (f) and MYC4 (g) protein stability in *35S:MYC3-GFP* and *cop1-4 35S:MYC3-GFP* seedlings grown in DD. MYC3-GFP/MYC4-GFP protein stability is elevated in MG132-treated seedlings in the wild-type background, similar to mock in the *cop1-4* background. (h) The MYC3 (left panel) and MYC4 (right panel) protein stability protein stability in *35S:MYC3-GFP, cop1-4 35S:MYC3-GFP, 35S:MYC4-GFP* and *cop1-4 35S:MYC4-GFP* seedlings grown in WL LD. In the presence of MG132, MYC3 protein gets stabilized, comparable to the DMSO-treated (mock) *cop1-4 35S:MYC3-GFP* seedlings. MYC4 protein stability was not altered in the MG132-treated samples compared to the mock, which is also similar in the *cop1-4* mutant background, suggesting COP1 does not control MYC4 protein stability in light. In (a-e), the total protein was extracted and subjected to immunoblot analysis using an anti-GFP antibody. Actin was used as a loading control for normalization in all the blots. Relative protein levels shown underneath each immunoblot were calculated using ImageJ software. The experiments were repeated two times, and similar results were obtained.

Further, to know if COP1-mediated degradation of MYC3 and MYC4 is through the 26S proteasomal pathway, we grew *35S:MYC3-GFP*, *cop1-4 35S:MYC3-GFP*, *35S:MYC4-GFP* and *cop1-435S:MYC4-GFP* seedlings in DD for five days, and on the sixth day, they were treated with either DMSO (mock) or MG132 for 16 h, and total protein was extracted and carried out immunoblotting. Our data suggest that in the MG132-treated samples, the MYC3-GFP was stable, but not in the mock (Figure 8f), as seen above. Similarly, MYC4-GFP was more stable in the MG132-treated samples than the mock in the wild-type background (Figure 8g); however, in the *cop1-4* mutant background, the stability of MYC4-GFP was comparable between MG132-treated samples and mock (Figure 8g). In WL-grown seedlings under LD, MG132 treatment slightly stabilized MYC3-GFP in the wildtype background, while in the *cop1-4* mutants, its stability was comparable to the mock (Figure 8h, left panel). The MYC4-GFP protein stability was comparable to the mock in the wild-type background in WL-SD (Figure 8h, right panel). In the *cop1-4* mutant background, MYC4-GFP protein stability in the MG132 samples was comparable to the mock, suggesting that COP1 probably doesn’t degrade MYC4 protein in light. These results reveal that COP1 targets MYC3 and MYC4 for degradation through the 26S proteasomal pathway in dark and light conditions.

### COP1 physically interact and ubiquitinates MYC3 and MYC4 before degradation

To understand in more detail whether COP1-mediated degradation of MYC3 and MYC4 is through direct physical interaction, we performed an in vivo co-immunoprecipitation assay from *35S:MYC3-GFP* and *35S:MYC4-GFP* transgenic seedlings grown in DD for five days and treated with MG132 for 16 h. Our immunoblot data suggest that when COP1 was co-immunoprecipitated using an anti-COP1 antibody in the immunoprecipitated complex, we could detect MYC3-GFP (Figure 9a) and MYC4-GFP (Figure 9b) using an anti-GFP antibody but not in the Col-0 (Figure 9a, b), which was used as a negative control. These results were further confirmed by BiFC assay in the onion epidermal cells. When cYFP-COP1 was co-infiltrated with nYFP-MYC3 or nYFP-MYC4, strong YFP fluorescence was detected in the nucleus (colour changed to green), as shown by the DAPI stain (Figure 9c). However, when cYFP/nYFP-MYC3, cYFP/nYFP-MYC4, cYFP-COP1/nYFP, and nYFP/cYFP empty vector combinations were co-infiltrated, no fluorescence was detected (Figure 9c).

**Figure 9.**
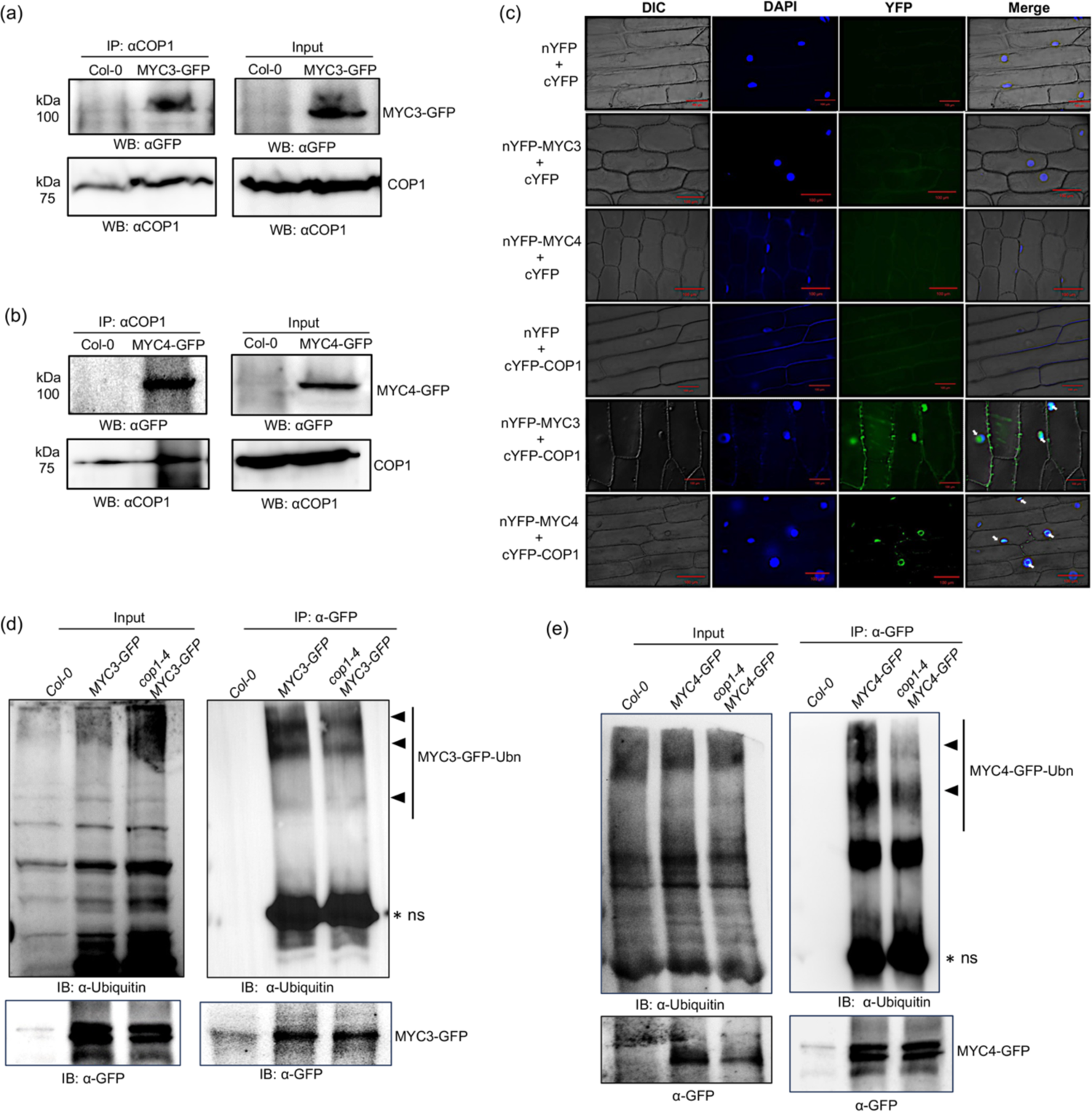
COP1 ubiquitinates and degrades MYC3 and MYC4, probably through direct physical interaction. (a, b) The co-immunoprecipitation assays reveal that COP1 physically interacts with MYC3 (a) and MYC4 (b). Five-day-old seedlings grown in DD conditions were treated with the proteasomal inhibitor MG132 for 16 h, tissue was harvested, and protein was extracted. Total protein was used for immunoprecipitation using anti-COP1 antibody. The immunoprecipitated protein complex was analyzed for MYC3-GFP and MYC4-GFP using anti-GFP antibodies. (c) BiFC assay for physical interaction between MYC3/MYC4 and COP1. All the constructs containing nYFP and cYFP were co-transformed into onion epidermal cells. The positive interactions were observed in nYFP-MYC3/cYFP-COP1 and nYFP-MYC4/cYFP-COP1 but in the negative control combinations. The left panel image shows the bright field image (DIC), the middle panel shows the nucleus staining by DAPI, and the next panel shows the YFP channel produced by the reconstruction of YFP. The right panel image shows the merged image. The Arrows indicate the reconstituted YFP signal in the nuclei. (d, e) In vivo ubiquitination assays show that COP1 ubiquitinates MYC3 (d) and MYC4 (e) in the dark as the ubiquitinated MYC3 and MYC4 levels decreased in the *cop1-4* mutant. Five-days-old etiolated *35S:MYC3-GFP* and *35S:MYC4-GFP* transgenic seedlings were treated with 50 µM MG132 in liquid MS for 16 h, followed by immunoprecipitation with magnetic beads conjugated with anti-GFP antibody. Input and IP samples were detected using anti-ubiquitin (upper panel) and anti-GFP (lower panel) antibodies.

Proteasomal degradation of substrates by COP1 requires ubiquitination of its target (Osterlund *et al*., 2000a, Holm *et al*., 2002, Seo *et al*., 2004, Kahle *et al*., 2020). As COP1 physically interacts with MYC3 and MYC4 and degrades them, we were curious to know if COP1 ubiquitinates MYC3 and MYC4 before degradation. To check this, we used *35S:MYC3-GFP*, *cop1-4 35S:MYC3-GFP*, *35S:MYC4-GFP* and *cop1-4 35S:MYC4-GFP* transgenic lines grown in DD for five days and treatment with MG132 for 16h, followed by protein extraction and immunoprecipitation with αGFP antibody. When we analyzed these immunoprecipitated samples using an anti-ubiquitin (αUbn) antibody, we could detect the ubiquitinated (Ubn) form of MYC3 and MYC4 proteins (Figure 9d, e), as both the proteins showed increased molecular weight (≥110 kDa) as compared to the non-ubiquitinated form (∼110 kDa) (Figure 9d, e). Moreover, the stability of the ubiquitinated versions of MYC3-GFP and MYC4-GFP was reduced in the *cop1-4* mutant comparison to the wild-type background (Figure 9d, e). Analysis of immunoprecipitated complex using αGFP antibody showed the protein stability of MYC3-GFP and MYC4-GFP was comparable between WT and the *cop1-4* mutant backgrounds (Figure 9d, e). These results confirm that COP1 ubiquitinates and degrades MYC3 and MYC4 in the dark to optimize seedling skotomorphogenic growth.

## Discussion

MYC transcription factors emerged as major regulators of growth and development in addition to their role in defense responses to herbivory and plant pathogens. Among MYCs, MYC2 is a dominant player regulating plant defense responses, while MYC3 and MYC4 have a redundant role (Niu *et al*., 2011, Ortigosa *et al*., 2020). Specifically, the role of MYC2 in light-mediated seedling development has been well established (Yadav *et al*., 2005, Gangappa *et al*., 2010, Gangappa *et al*., 2013b, Sethi *et al*., 2014, Chakraborty *et al*., 2019, Ortigosa *et al*., 2020, Srivastava *et al*., 2022). However, the role of MYC3 and MYC4 and their interaction with MYC2 in the seedling photomorphogenesis was unknown. This study demonstrates the coordinated interaction between MYC2/MYC3/MYC4 in regulating seedling photomorphogenic growth. Through genetic and molecular analyses, we have uncovered the multifaceted roles of MYC3 and MYC4 in modulating various aspects of light-mediated growth and development, shedding light on their interplay with key regulatory factors such as COP1 and HY5 (Figure 10).

**Figure 10.**
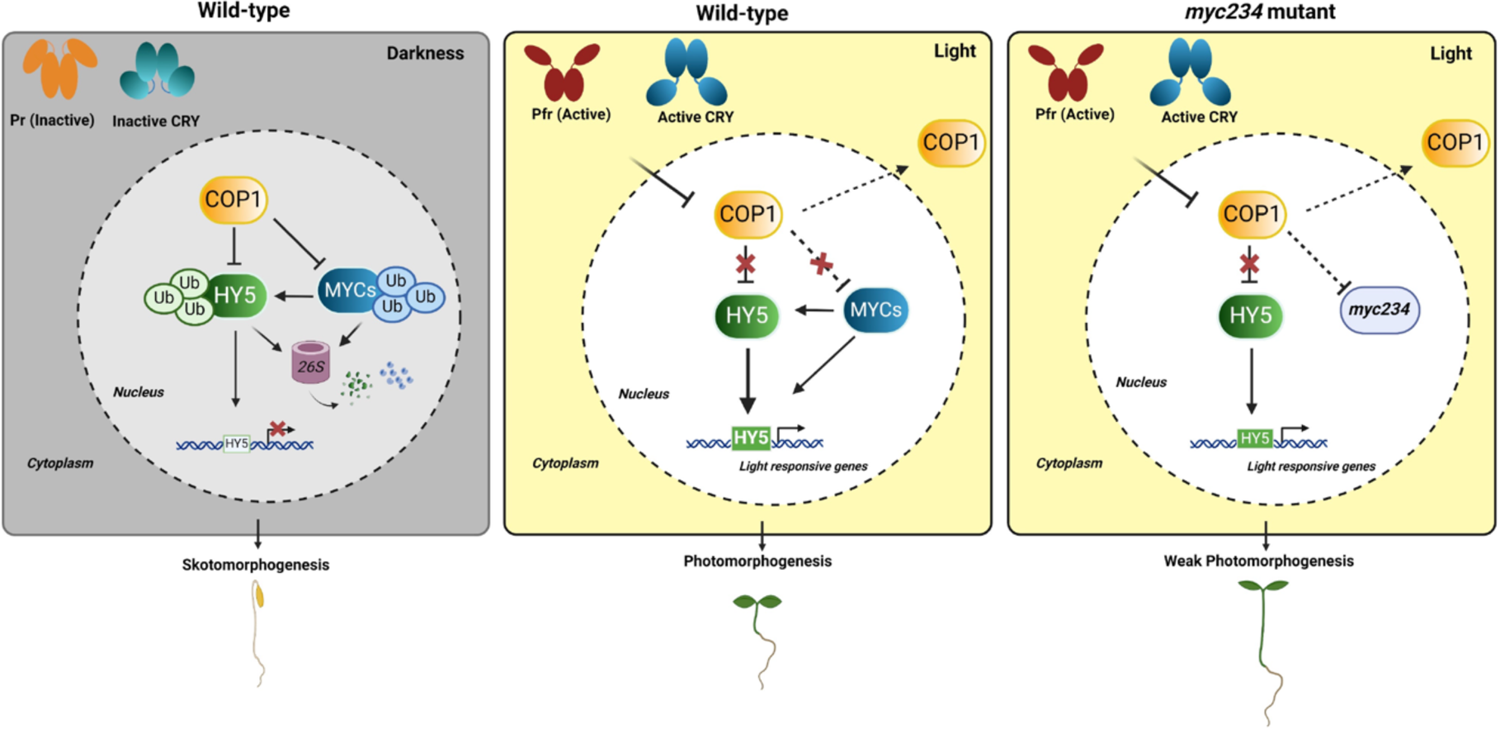
Working model of MYC transcription factors in the regulation of seedling photomorphogenesis. In the dark, COP1 targets HY5 for degradation to promote skotomorphogenesis. At the same time, COP1 also degrades MYCs for optimal skotomorphogenic growth. In the light, photoreceptor-mediated inhibition of COP1 and partial depletion of COP1 from the nucleus due to cytosol result in enhanced HY5 accumulation. Also, enhanced accumulation of MYCs leads to increased HY5 activity, which leads to optimal photomorphogenic growth. It is also likely that MYCs together promote photomorphogenesis independent of HY5. The absence of all the MYCs (*myc234* triple mutant) reduces HY5 activity, likely due to reduced *HY5* transcript accumulation, resulting in weak photomorphogenic growth. In summary, MYC2/3/4 together promotes photomorphogenesis by enhancing HY5 function, while individually, they act as negative regulators of photomorphogenesis.

Our study unravels MYC4 as a negative regulator of seedling photomorphogenesis in a wavelength-independent manner. While MYC3 is found to have a minor role on its own, together with MYC4, it strongly enhances the photomorphogenic growth of seedlings as the double mutant of *myc3myc4* displayed significantly longer hypocotyls than the *myc3* and *myc4* (Figure 1). These results suggest that MYC3 and MYC4 have independent and interdependent functions and display unequal redundancy in regulating seedling hypocotyl growth type (Figure 1). Unlike their role in hypocotyl growth, MYC3 and MYC4 together probably play a negative role in light-mediated cotyledon growth, as the *myc3myc4* double mutants had bigger cotyledons than the *myc3*, which has moderately bigger cotyledons than wild-type. The lack of cotyledon phenotype in the *myc4* mutant suggests that MYC4 functions partially redundantly with MYC3 in inhibiting cotyledon expansion in response to light. Consistent with the hypocotyl growth phenotype, MYC transcription factors differentially regulate the anthocyanin and chlorophyll accumulation and the expression of light-inducible genes, as the anthocyanin, chlorophyll content, and gene expression levels in the triple mutants were very low compared to the single mutants, wherein it was either more or similar to wild-type (Figure 2).

Unequal genetic redundancy is a common phenomenon in Arabidopsis (Laubinger *et al*., 2004, Dohmann *et al*., 2005, Sibout *et al*., 2006)and likely in crops and other models (Briggs *et al*., 2006). For example, MYC2 shows unequal genetic redundancy with a bZIP transcription factor GBF1 in regulating hypocotyl growth (Maurya *et al*., 2015). Both *myc2* and *gbf1* have shorter hypocotyl than the wild-type (Yadav *et al*., 2005, Mallappa *et al*., 2006); in the *myc2gbf1* double mutant, instead of enhancing the short hypocotyl phenotype, they suppressed their short hypocotyl phenotype and had hypocotyl length similar to the wild-type (Maurya *et al*., 2015). Similarly, *spa1* mutants have a reduced hypocotyl length, while spa2 mutants show a hypocotyl length similar to that of wild-type. However, the *spa1spa2* double mutants show shorter hypocotyls than the *spa1,* suggesting that SPA1 and SPA2 show unequal redundancy in controlling hypocotyl growth (Laubinger *et al*., 2004). Similarly, the two closely related bZIP transcription factors, which are key regulators of photomorphogenic growth, showed reduced root system growth phenotype in the *hy5hyh* double mutants than in the single mutants, wherein the root system growth is enhanced (Sibout *et al*., 2006). The COP9 signalosome (CSN) components, CSN5A and CSN5B, also have similar genetic interactions for the regulation of seedling photomorphogenesis and lateral root development, as the *csn5b* mutant does not show any phenotypic changes further enhance the *csn5a* mutant phenotypes (Dohmann *et al*., 2005). This evidence supports our observations that the mode of MYCs function varies when they function alone or with other MYC family members, suggesting that they probably have independent and interdependent functions, highlighting the existence of unequal genetic redundancies among MYC transcription factors underlying photomorphogenic responses.

Consistent with the *myc234* triple mutant phenotypes, such as long hypocotyl, reduced anthocyanin and chlorophyll accumulation, our results also revealed reduced HY5 protein stability in the *myc234* triple mutants under WL and BL. HY5 is a key activator of genes involved in anthocyanin biosynthesis (Ang *et al*., 1998, Chattopadhyay *et al*., 1998a). The reduced expression of these genes in *myc234* further supports our hypothesis that MYCs promote seedling photomorphogenesis by probably enhancing HY5 signaling by stabilizing HY5 protein. However, unlike in BL, HY5 protein stability was elevated in the *myc234* triple mutant in RL and FRL conditions, highlighting that MYCs differentially regulate HY5 protein stability in a wavelength-dependent manner. Our investigation into the genetic interaction between MYC3/MYC4 and HY5 revealed a nuanced regulatory mechanism governing hypocotyl growth. While HY5 is a well-established positive regulator of the photomorphogenesis (Ang and Deng, 1994, Oyama *et al*., 1997, Ang *et al*., 1998, Chattopadhyay *et al*., 1998b, Gangappa and Botto, 2016), our results demonstrate that MYC3 and MYC4 genetically interact with HY5, potentially acting as negative regulators to suppress its function. However, the ability to suppress *hy5* mutant hypocotyl length is slightly relieved in the presence of *myc3myc4* mutations. The role of MYC3 and MYC4 in suppressing HY5 functions is also consistent with MYC2 suppressing HY5 function to inhibit seedling photomorphogenic growth (Chakraborty *et al*., 2019). This antagonistic relationship underscores the complexity of regulatory networks controlling seedling development and emphasizes the role of MYC transcription factors in modulating the activity of key photomorphogenic regulators (Ang and Deng, 1994, Ang *et al*., 1998).

On the one hand, our genetic and molecular analysis revealed that MYC3 and MYC4 are probably acting to promote HY5 functions, as revealed by the hypocotyl length phenotype. On the other hand, COP1 targets MYC3 and MYC4 for ubiquitination and subsequent proteasomal degradation, thereby modulating their abundance in response to light stimuli. COP1 exerts a stricter control on MYC3 under dark and light conditions, as MYC3-GFP is detectable only in the background of the *cop1-4* mutation or when the wild-type seedlings were treated with the proteasomal inhibitor MG132 (Figure 8). The *myc3* mutant alone does not display any visible changes in the hypocotyl length, while in the *myc2* and *myc4 background,* it modulates their phenotype. MYC3 is likely playing a crucial role in regulating the functions of MYC2 and MYC4, and therefore, COP1 plays a role in this modulation by maintaining very low MYC3 protein levels. While MYC4 is also targeted by COP1 for ubiquitination and degradation, unlike MYC3, MYC4 protein remains present in dark and light conditions, albeit at a lower level. In response to MG132 treatment, MYC4 protein stability increases to a level similar to mock-treated *cop1-4* seedlings in the dark (Figure 8). Consistent with increased MYC3 and MYC4 protein stability in the dark upon MG132 treatment, the ubiquitinated form of MYC3-GFP and MYC4-GFP was visibly reduced in the *cop1-4* mutant compared to wild-type seedlings, suggesting that the absence of COP1 results in lack of physical interaction and reduced ubiquitination and enhanced MYC3/MYC4 protein stability (Figure 9). Like MYC3 and MYC4, MYC2 was also shown to be degraded by COP1 both under constant dark and diurnal light/dark conditions (Chico *et al*., 2014).

COP1 is the central regulator of light signaling, which regulates the activity of transcription factors through degradation in fine-tuning the balance between skotomorphogenic and photomorphogenic growth. Our findings are consistent with previous reports highlighting the pivotal role of COP1 in orchestrating light-mediated developmental processes by regulating the protein stability of transcription factors (Osterlund *et al*., 2000b, Holm *et al*., 2002, Saijo *et al*., 2003, Seo *et al*., 2004). Together, our study provides a mechanistic insight into the complex regulatory mechanism through which MYC transcription factors regulate seedling photomorphogenesis (Figure 10). While COP1 controls MYCs function by regulating their protein degradation, MYCs, in turn, control photomorphogenic responses by controlling HY5 protein activity or its function, and also plausibly by independent of HY5. The reduced protein stability of HY5 in the *myc234* triple mutants is likely due to reduced *HY5* transcription (Figure 2), consistent with earlier reports (Ortigosa *et al*., 2020). By elucidating the roles of MYC3 and MYC4 in Arabidopsis seedling development, our findings contribute to growing evidence of complex genetic interactions among the members of the same family or between different transcription factors. These interactions are necessary for fine-tuning the signaling to control various aspects of plant growth and development in response to changing light environments.

## Material and Methods

### Plant material and growth conditions

This study uses Arabidopsis thaliana Columbia ecotype (Col-0) as wild-type plant material. The *myc3* (GK_445B11) and *myc4* (GK_491E10) have been ordered from the Nottingham Arabidopsis Stock Centre (NASC). The *myc2* (*jin1-2*) allele used in this study is described elsewhere (Lorenzo *et al*., 2004). The *cop1-4* (McNellis et al., 1994) and *hy5-215* (Oyama *et al*., 1997) are described elsewhere. The *35S:MYC3-GFP* and *35S:MYC4-GFP* are described elsewhere (Fernandez-Calvo *et al*., 2011). All the mutants used in this study are in Col-0 background.

Seeds were surface sterilized (70% ethanol + 0.05% Triton X-100) and plated on Murashige and Skoog agar medium (MS plates) with 1% sucrose. The plates were then cold-treated in the dark for stratification for four days and transferred to light chambers maintained at 22°C with the required wavelength and intensity. The 100-120 mmol m^-2^ s^-1^ light intensity is used to grow seedlings for all the experiments. Experiments are performed under specified short-day conditions (8-h-light/16-h-dark) or long-day (16-h-light/8-h-dark) conditions.

### Generation of double and triple mutants

For the generation of the *myc3myc4* (*myc34*) double-mutant, homozygous *myc3* mutant plants were crossed with *myc4* homozygous lines. The F2 seedlings were selected in MS with sulfadiazine antibiotic plates. F2 plants were then tested by genomic PCR for both *myc3* and *myc4* loci. F3 progeny that is homozygous for *myc3* and *myc4* mutation is further tested by genotypic PCR and is designated as *myc34* double-mutants. In the same process, single mutant *myc3* and *myc4* are crossed with the *hy5* homozygous mutant, respectively. The *hy5* mutant seedlings were selected in the F2 population. F2 plants were then individually tested by genomic PCR for the myc3 and myc4 locus. F3 progeny that is homozygous for *myc3* mutation is further tested by genotypic PCR and designated as *myc3hy5* double-mutants, and those are homozygous for *myc4* mutation and PCR tested, designated as *myc4hy5* double-mutants. In the same way, *myc3cop1-4, myc4cop1-4, myc34cop1-4* mutants were generated through genetic crossing. In all the cases, homozygous seeds were confirmed by PCR screening and bulked seeds for the experiments.

Transgenic seedlings overexpressing MYC3 and MYC4 in *cop1-4* mutants were generated by genetic crosses using *cop1-4* single mutants as females and MYC3 and MYC4 transgenic lines as males in each of the individual crosses. Seedlings with *cop1* mutant phenotype were selected in F2 populations, and the overexpression of *MYC3-OE* and *MYC4-OE* transgene in *cop1-4* mutant was confirmed by western blot (using anti-GFP antibodies). In the same way, *MYC3-OE* and *MYC4-OE* transgenic lines were made in the background of the *hy5* mutant. Several homozygous lines were reconfirmed in F3 generation and were used for further studies.

### Easy plant DNA extraction for genotyping the mutants

Small leaf discs were collected in 40 µl of 0.25M NaOH in sterile Eppendorfs. Samples are incubated in a water-bath for 30 sec. Samples are neutralized by adding 40 µl of 0.25 M HCl and 20µl of 0.5 M Tris-HCl of pH 8.0 containing 0.25% (V/V) NP-40 or triton-X 100. The samples were boiled again for 2 mins and stored at 4°C.

### Hypocotyl length measurement

Six-day-old seedlings grown as specified above are used for hypocotyl measurements. At least 20-25 seedlings were aligned on a 1% agar plate containing 1% charcoal and imaged using a digital camera, and hypocotyl lengths were measured using NIH ImageJ software.

### RNA extraction and gene expression analysis by qPCR

Six-day-old dark/light-grown seedlings were used for the total RNA isolation. Approximately 100 mg of plant seedlings/other tissues were powdered using liquid nitrogen before extracting total RNA using RNeasy® Plant Mini Kit (QIAGEN) with on-column DNase I digestion according to the manufacturer’s instructions. The integrity of RNA was quantified by Nanodrop. Approximately 2.0 µg of total RNA was converted into cDNA using a verso cDNA synthesis kit (Thermo-Fisher scientific) and oligo dT following the manufacturer’s instructions. 2.0 µL of 1:20 diluted cDNA was used for qPCR using PowerUp SYBR Green Master Mix. qPCR experiments were performed in QuantStudio™ 5 Real-Time PCR System. *EF1α* (AT5G60390) was used as an internal control for normalization. Details of the oligonucleotide primers used are provided in Table S1.

### Anthocyanin level measurement for Arabidopsis seedlings

About 20–30 seedlings are taken into a microcentrifuge tube and weighed (around 35-50mg); 400 µl of extraction solution (1% HCL in Methanol) is added and vortexed very well and kept at 4°dark conditions overnight. The next day, the seedlings were crushed, and 200µl of sterile water and 200 µl of chloroform were added. The debris was removed by centrifugation (10 minutes, 13000 rpm), and the supernatant was collected into a fresh microcentrifuge tube. Then, spectrophotometric estimation is carried out by taking readings at 530 nm and 657 nm wavelengths. The total Anthocyanin content is calculated with the help of the following formula: (A530-0.33A657)/gm of tissue.

### Chlorophyll estimation for Arabidopsis seedlings

Approximately 30-40 six-day-old seedlings were collected, weighed, frozen in liquid nitrogen, and crushed to a fine powder for the chlorophyll measurements. After extracting the total chlorophyll into 1 mL of 80% acetone and centrifuging for 10 minutes at 13000 rpm, the amounts of chlorophyll a and b were determined using MacKinney’s specific absorption coefficients, with chlorophyll A equaling 12.7(A663) - 2.69(A645) and chlorophyll B equaling 22.9(A645) - 4.48. (A663). The chlorophyll content is expressed as Total chlorophyll/gram of seedling tissue.

### Protein extraction and immunoblotting analysis

Approximately 100 mg of tissue was harvested in a microcentrifuge tube, snap frozen in liquid nitrogen and ground in 200 µl of protein extraction buffer (50 mM Tris HCl pH 8.0, 150 mM NaCl, 10% glycerol, 5mM DTT, 1% (v/v) Protease Inhibitor Cocktail, 1% NP40, 0.5mM PMSF). The protein extract was centrifuged at 10,000 rpm for 15 min to pellet down the debris at 4°C. The supernatant was then transferred to a fresh tube, and an aliquot of 3-5 µl was taken out in a separate tube to estimate protein by Bradford assay. The protein samples (1µg/µl) were boiled for 5-10 min at 100°C.

The denatured protein samples were loaded onto the gel. After separating the protein samples on SDS-PAGE gel, they were transferred to PVDF membrane at 90V for 1 h in transfer buffer (Tris 48 mM, Glycine 39 mM, 20% methanol pH 9.2) in wet transfer method in cold room. The membrane was stained with Ponceau-S to confirm the protein transfer and then washed with sterile MQ water. The membrane was then incubated on a rotary shaker for 2 h in 10 ml blocking buffer (5% non-fat dry milk in TBS and 0.05% Tween-20) at room temperature. The blocking reagent was removed, and the affinity-purified primary antibody diluted (1:250 to 1:10,000) in 10 ml TBS with 0.05% Tween-20 was added and incubated overnight with shaking in a cold room. The membrane was then washed thrice with 10 ml of wash buffer (TBS and 0.05% Tween-20) for 5 min each. The secondary antibody goat anti-mouse (Invitrogen, 31430), goat anti-rabbit (Abcam, ab205718) HRP conjugated was diluted (1:5,000 to 10,000) in 10ml TBS with 0.05% Tween-20 was added and incubated for 1 h with shaking at room temperature. The membrane was washed thrice with 10 ml of wash buffer at room temperature. Western blot was performed using the Super Signal West Femto chemiluminescent substrate kit (Pierce) and following the manufacturer’s instructions. A substrate working solution was prepared by mixing peroxide solution and Luminol/enhancer solution in a 1:1 ratio, and the blot was incubated in that working solution for 5 min in the dark. The blot was then removed from the working solution and observed in Chemi-Doc at different times depending on signal strength.

### Bi-molecular fluorescence complementation assay (BiFC assay)

For the BiFC studies, full-length CDS of MYC3 and MYC4 was cloned in pENTR/D-TOPO (Invitrogen) to generate *pENTR-MYC3* and *pENTR-MYC4*. Then, LR Clonase II enzyme mix was used to generate destination construct pSPYNE vector to obtain MYC3-YFPN-ter (MYC3-nYFP) and MYC4-YFPN-ter (MYC4-nYFP), respectively. Similarly, full-length CDS of HY5 was cloned in pSPYCE to obtain HY5-YFPC-ter (HY5-cYFP), and full-length CDS of COP1 was cloned in pSPYCE to obtain COP1-YFPC-ter (COP1-cYFP). The constructs and *p19* plasmid were transformed in *Agrobacterium* (GV3101 strain). A single colony of the transformed Agrobacterium was inoculated in 5 ml of YEM media in each case and incubated at 28°C at 200 rpm overnight. The cultures were then centrifuged for 15 min at 4000 g, and the pellets were resuspended in 1 ml of AS medium (100 mM MES-KOH pH 5.6, 100 mM CaCl2, 100 µm acetosyringone). The cell density was diluted with AS medium to OD600 ∼ 0.7–0.8. The different Agrobacterium strains harbouring the respective constructs were mixed in a ratio of 1:1:0.5 (total volume of 3 ml per leaf) and were allowed to stand for 2–4 h at room temperature. About 200 ml agroinfiltration liquid was slowly injected into the interfaces between the adaxial epidermis and mesophyll of onion bulb scales by using a plastic syringe with a needle, which resulted in an agroinfiltration bubble at the injection spot filled with infiltration liquid occupying about 1 cm^2^ area of the epidermis. The fluorescence in the epidermal cell layer expressing the fusion protein was observed after three days of infiltration under the confocal microscope.

### Co-Immunoprecipitation assay

The Col-0, *35S:MYC3-GFP* (MG132 treated) and *35S:MYC4-GFP* overexpression lines grown either in dark or light for six days were used for in-vivo protein-protein interaction assay. Approximately 500 μg of total protein of each line were extracted in the buffer containing 50 mM Tris-Cl, pH 8.0, 150 mM NaCl, 10% glycerol, 5 mM DTT, 1% (v/v) protease inhibitor cocktail, 1% NP40, 0.5 mM PMSF and 0.5 mM β-Mercaptoethanol. The Dynabeads protein G-magnetic beads (Invitrogen) and 2.5 µl of anti-GFP polyclonal antibody (Abcam, ab290) were incubated with Ab binding and washing buffer (provided in the kit) overnight at 4°C in the rocker. The next day, the Eppendorf tubes were placed onto a magnetic stand, and the supernatant was discarded. The magnetic bead-Ab-protein complex was resuspended in Ab-binding and washing buffer by gentle pipetting. The extracted protein samples were mixed into the buffer and kept overnight at 4°C. The next day, magnetic beads were separated using a magnetic stand and washed thrice with a washing buffer (provided in the kit). The eluate was collected and boiled at 70°C for 10 min and then run in SDS-PAGE. Both input and IP were analyzed by probing with an anti-HY5 antibody (Agrisera, AS12 1867) and an anti-COP1 antibody (Agrisera, AS20 4399). For checking the protein-protein interaction of MYC3 and MYC4 with COP1, 2.5 µl of anti-COP1 antibody was used for IP reactions, and immunoprecipitated samples were analysed for the presence of MYC3 or MYC4 using anti-GFP antibody.

### Proteasomal degradation and Ubiquitination assay

For this experiment, five-day-old dark-grown seedlings of Col-0, *MYC3-GFP, cop1-4 MYC3-GFP, MYC4-GFP* and *cop1-4 MYC4-GFP* were treated with MG132 (50 µM *Z-Leu-Leu-Leu-al,* Sigma) or mock (DMSO) in liquid MS for 16 h. The seedlings were harvested, and total protein was extracted in a protein extraction buffer. 50 µM MG132 was added to the protein extraction buffer during protein extraction from the treated sample. For the mock samples, DMSO was used in the protein extraction buffer. The boiled protein samples were loaded into SDS-PAGE gel, and a western blot was performed. The co-immunoprecipitation assay was performed by using Dynabeads protein G-magnetic beads (Invitrogen). For IP reactions, an anti-GFP monoclonal antibody (Takara, 632592) was used. All eluted immunocomplex and input samples were boiled at 70°C for 10 min before immunoblotting. The proteins were analysed by immunoblotting with anti-ubiquitin (Invitrogen, eBioP4D1 (P4D1), and anti-GFP (Takara, 632592) antibodies.

### Statistical analysis

For measuring hypocotyl length, at least 20-25 seedlings were used. Three independent biological replicates in combination with three technical replicates were used for gene expression analysis. All experiments were repeated at least thrice. The statistical significance between or among treatments and genotypes was determined based on a one-way analysis of variance or a two-way-analysis of variance test (ANOVA) according to the nature of the data. The error bar represents standard deviations.

## Author contributions

S.N.G. conceptualized and supervised the study. V.G. conducted most experiments, collected results, analysed the data, and prepared figures. S.D. performed gene expression analysis. S.N.G. wrote the manuscript with the help of V.G. All the authors have read and approved the final version.

## Acknowledgements

We thank all the members of the Gangappa laboratory for the helpful discussions. We also thank Dr Channakeshavaiah C. for the BIFC binary vectors, Prof. Shubho Chaudhuri for the GV3101 agrobacterium strain, and Prof. Roberto Solano for the *jin1-2*, *35S:MYC3-GFP* and *35S:MYC4-GFP* transgenic seeds. V.G. and S.D. acknowledge the University Grants Commission (UGC, Govt of India) and IISER Kolkata (Ministry of Education) for their doctoral fellowship. We acknowledge Mr Chirag Singhal for COP1 BIFC constructs; we also thank Prof. Sumana Annagiri (IISER Kolkata) for the microscopy and the BioRender for allowing us to draw our model.

## Data availability statement

All the data presented in this study can be found in the main manuscript and the supporting information.

