## Supplementary data for "Unequal genetic redundancies among MYC bHLH transcription factors underlie seedling photomorphogenesis in Arabidopsis"

### SUPPLEMENTAL INFORMATION

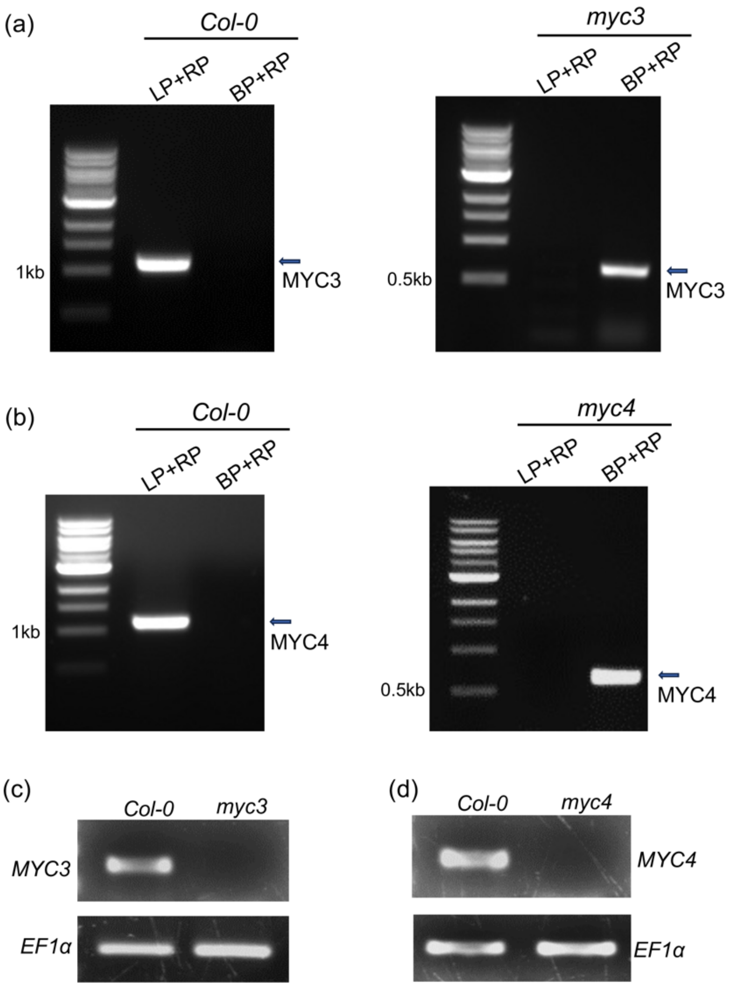


**Figure S1: Identification and molecular characterization of *myc3* and *myc4* T-DNA insertion mutant lines.**

(a, b) Genomic PCR analysis of *myc3* (a) and *myc4* (b) mutants with respective gene-specific and border primers to detect the presence or absence of T-DNA insertion in *MYC3* and *MYC4* loci. The forward/reverse primers corresponding to *MYC3* or *MYC4* genomic sequences flanking the T-DNA insertion and the left border primer of the T-DNA insertion corresponding to the sequence of the T-DNA fragment were designed for PCR analysis. The arrows indicate the expected band size for genomic fragment or genomic + T-DNA fragment.

(c) Semi-qRT PCR of MYC3 and MYC4 genes from cDNA from Col-0 for full-length transcript where bands were absent in myc3 and myc4 mutants. *EF1α* was used as equal RNA loading control.

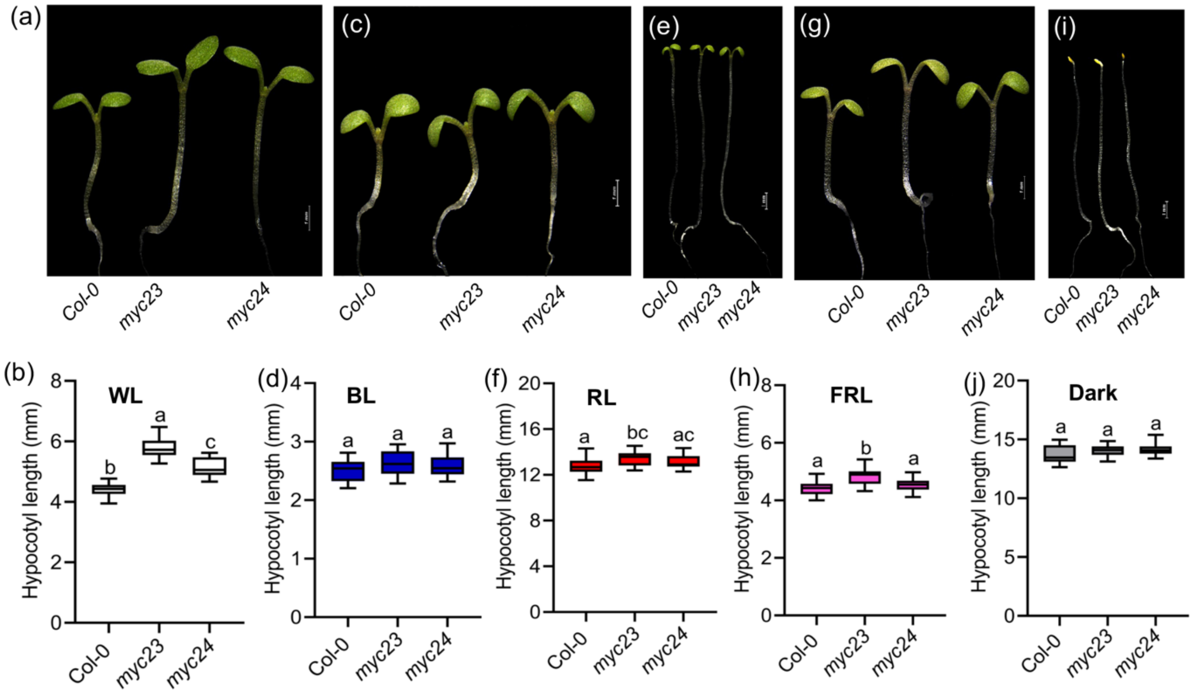


**Figure S2. MYC2 genetically interact with MYC3 and MYC4 to control growth in seedling photomorphogenic growth**

(a-h) Representative seedling images and hypocotyl length of six-day-old Col-0, *myc23* and *myc24* seedlings grown under SD in WL (a, b), BL (c, d), RL (e, f), FR (g, h) or in DD (i, j). Box-whisker plots represent mean±SD (n ≥25 seedlings). Different letters above the box plots indicate a significant difference (one-way ANOVA with Tukey's HSD test, $P < 0.05$), while the same letters indicate that the genotypes hypocotyl lengths do not differ significantly.

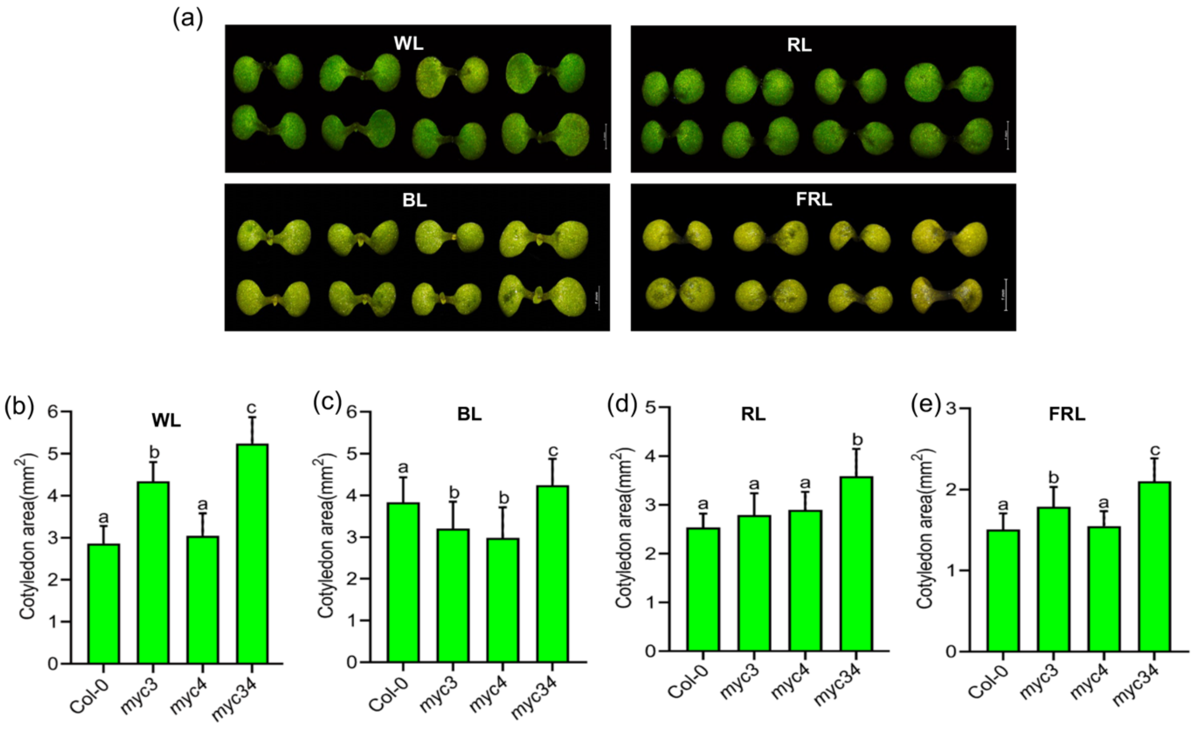


**Figure S3. MYC3/MYC4 interaction regulates cotyledon growth**

The cotyledons were excised from six-day-old seedlings and aligned on agarose media. The cotyledon area was measured using Image J software.

(a-e) Representative cotyledon images (a) and measured cotyledon area for six-day-old seedlings grown in WL (b), BL(c), RL(d) and FRL (e), respectively. Box-whisker plots represent mean ± SD (n ≥20 seedlings). Different letters in the box plots denote a significant difference (one-way ANOVA with Tukey's HSD test, $P < 0.05$).

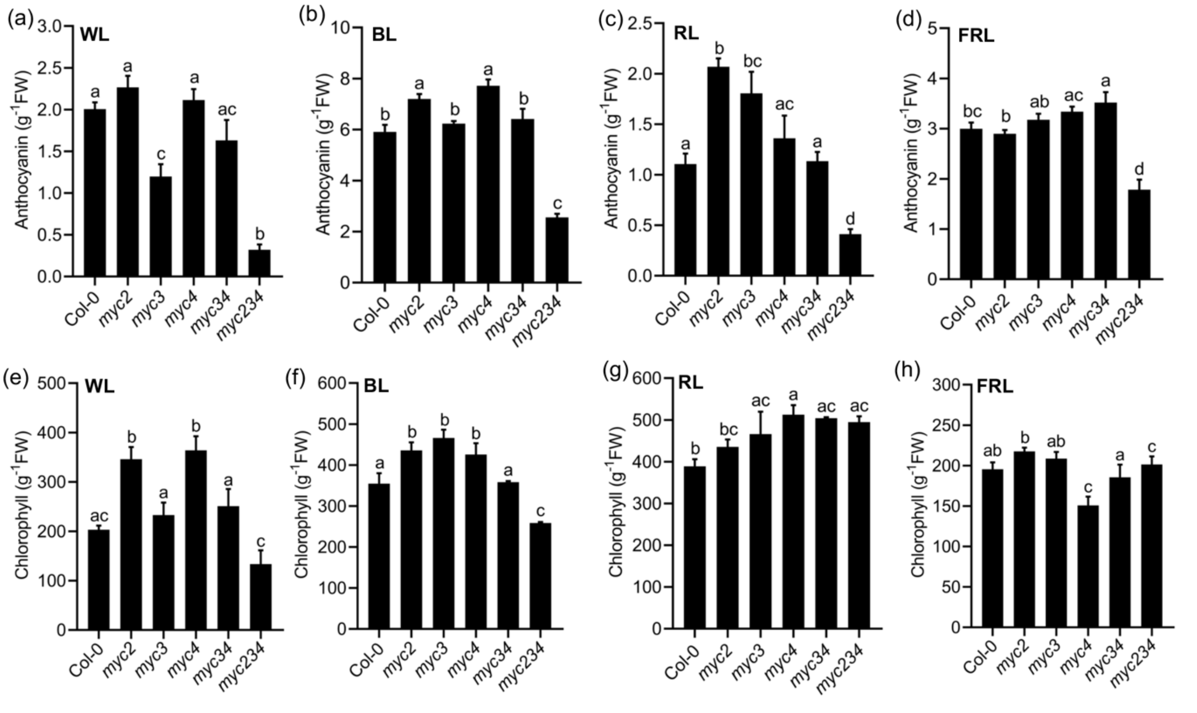


**Figure S4. MYC4 genetically interact with MYC3 and MYC2 to control anthocyanin and chlorophyll accumulation**

(a-d) Quantification of anthocyanin content in 22°C SD grown WL (a), BL (b), RL (c) and FRL (d) in six-day-old Col-0, *myc2, myc3, myc4*, *myc34* and *myc234* triple mutants.

(e-h) Chlorophyll content in six-day-old seedlings of Col-0, *myc2, myc3, myc4*, *myc34* and *myc234* triple mutants grown in WL (e), BL (f), RL (g), FRL (h) under SD photoperiod.

In panels (a-h), Different letters above the bar chart indicate a significant difference (one-way ANOVA with Tukey's HSD test, $P < 0.05$ ($n \geq 25$ seedlings). In contrast, the same letters show that the genotypes do not differ significantly.

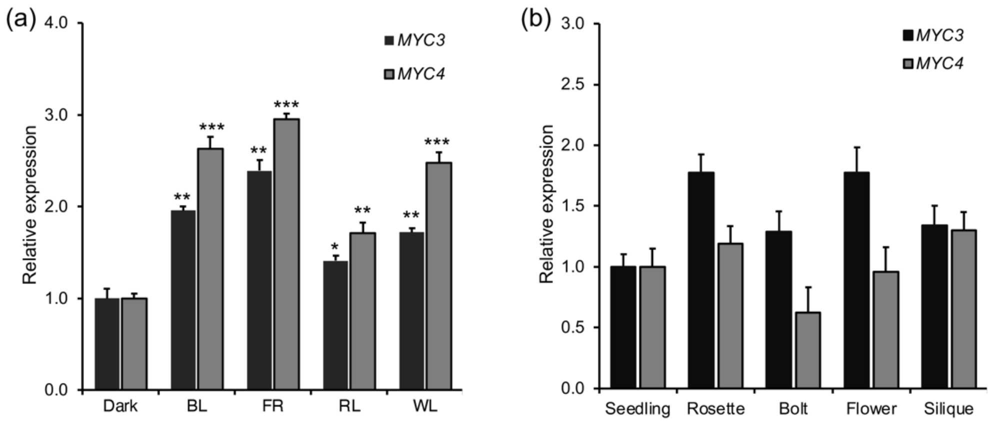


**Figure S5. *MYC3* and *MYC4* gene expression in seedlings and adult plant tissues.**

(a and b) Gene expression analysis of targets MYC3 and MYC4 in Col-0 under Dark and different light conditions in SD from six-day-old seedlings grown (a) and in different tissue types such seedlings, rosette, bolt, flower and siliques grown under LD condition (b) at 22°C. *EF1α* was used as housekeeping control. Asterisks denote statistically significant differences in different light conditions compared to dark as determined by Student's t-test (*P<0.05, **P<0.01, ****P*<0.001).

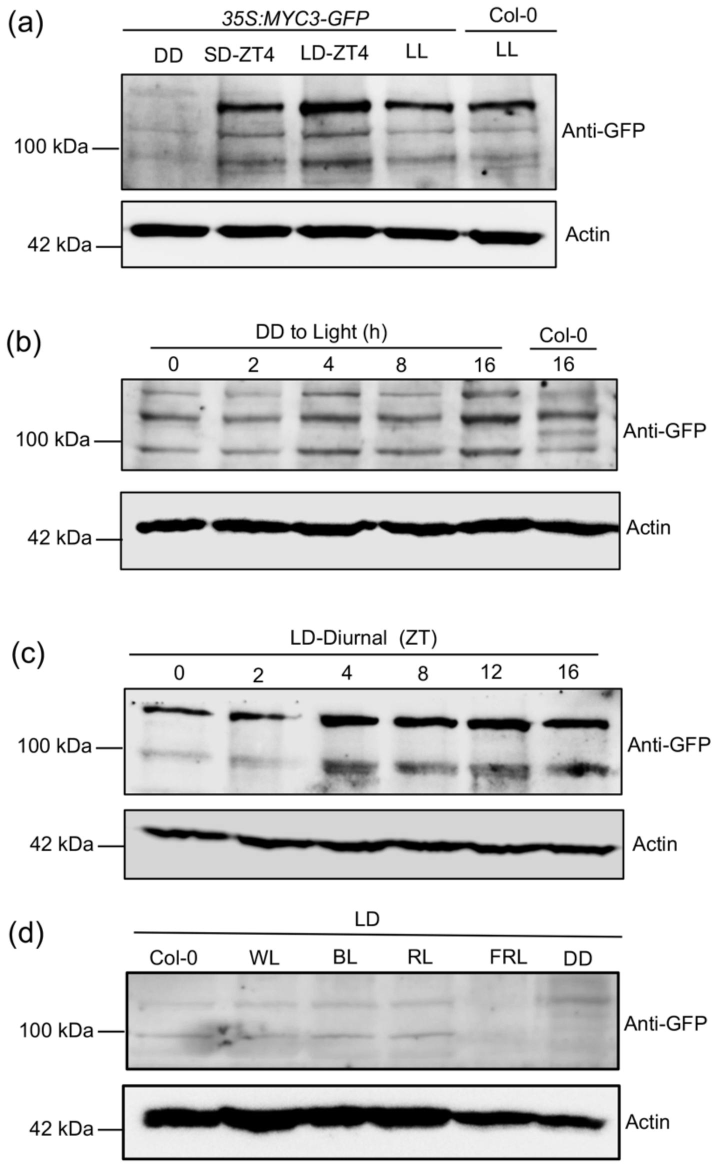


**Figure S6. Immunoblotting data shows that MYC3 protein is highly unstable in dark and light.**

(a-d) Immunoblot analysis of *35S:MYC3-GFP* transgenic line using anti-GFP antibody from constant dark and different photoperiods (a), in the dark to light shifted seedlings (b), under diurnal LD photoperiod (c), and in various light conditions such as DD, WL and monochromatic lights (d). Actin was used as a loading control. Col-0 was a negative control to track the band in the *35S:MYC3-GFP* line. Please note that the specific band corresponding to MYC3-GFP was observed in none of the conditions.

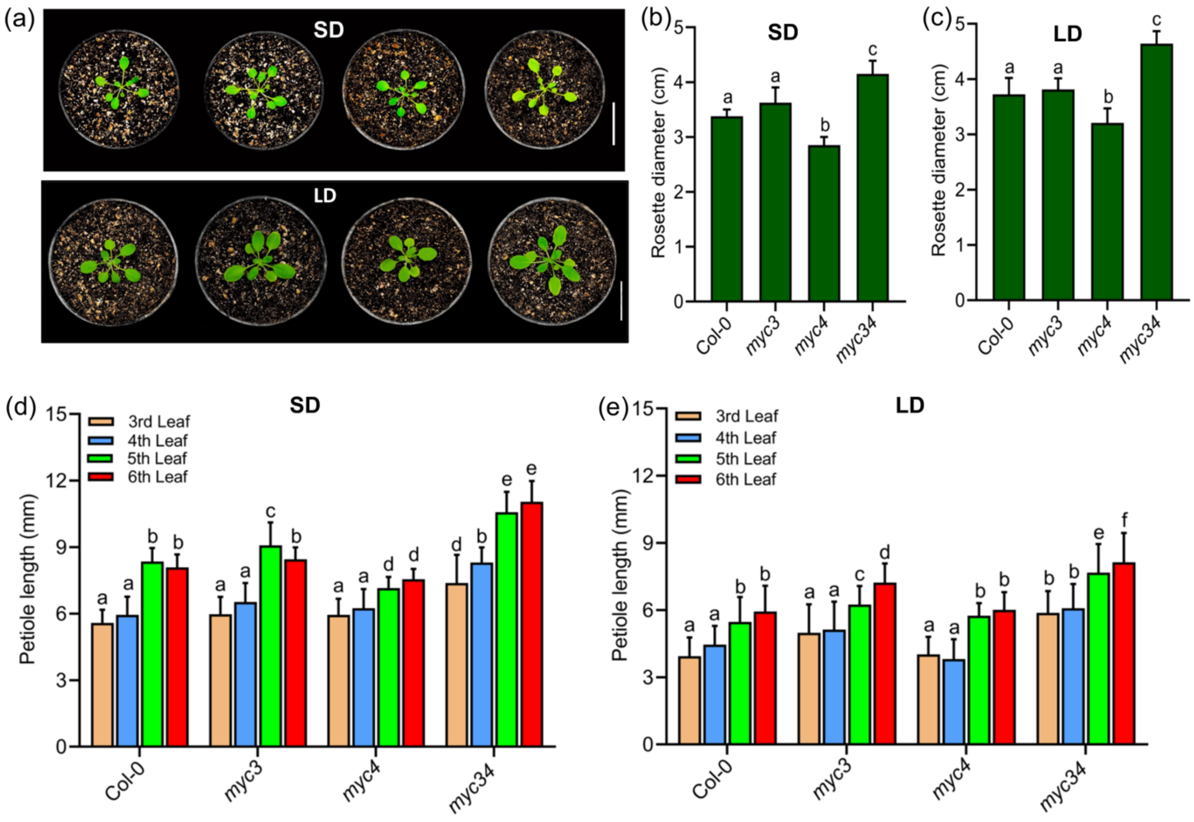


**Figure S7. MYC3/MYC4 are responsible for regulating petiole length and rosette phenotype.**

(a-c) Representative rosette images of three-week-old adult plants of Col-0, *myc3*, *myc4* and *myc34* genotypes grown under SD (top panel, a) and LD condition (bottom panel, a) and quantification of rosette diameter from SD (b) and LD (c) grown plants.

(d, e) Quantifying petiole lengths from three-week-old plants grown under SD (d) and LD (e) photoperiods. The 3$^{rd}$, 4$^{th}$, 5$^{th}$ and 6$^{th}$ leaves were excised, aligned on the agarose media, and photographed before being measured using ImageJ NIH software. Box-whisker plots represent mean ± SD (n ≥6 plants). Different letters in the box plots denote a significant difference (one-way ANOVA with Tukey's HSD test, $P < 0.05$).

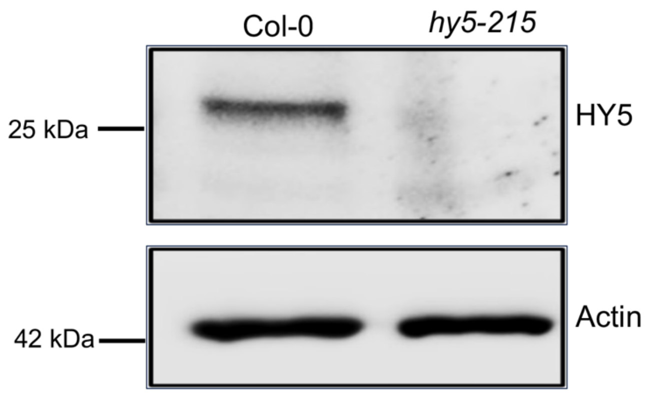


**Figure S8. The native anti-HY5 antibody specifically recognizes the HY5 protein.**

**(**a) Western blot analysis of Col-0 and *hy5-215* mutant grown under SD condition at 22°C. The native anti-HY5 antibody detected the HY5 protein in Col-0 but not in the *hy5-215* mutant. Actin was used as a loading control.

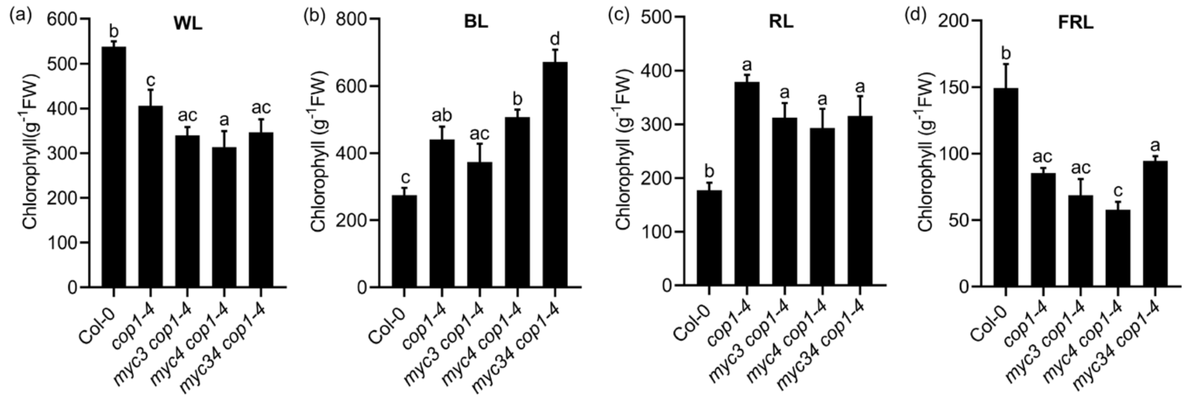


**Figure S9. The *myc3* and *myc4* mutations do not control the chlorophyll levels of the *cop1-4* mutant.**

(a-d) Accumulation of chlorophyll levels in six-day-old seedlings of the indicated genotypes grown in WL (a) and different monochromatic lights such as BL (b), RL (c) and FRL (d) and under SD photoperiod.

**Table S1. List of oligonucleotides used in this study.**

| Name | Oligo Sequence (5'…………3') | Purpose |
|---|---|---|
| **Primers used for genotyping PCR analysis** | | |
| *myc3*-LP | AGGCAAAACCCATTTACAACC | Genotyping |
| *myc3*-RP | TGAAGCAGAGAGGCAGAGAAG | Genotyping |
| *myc4*-LP | CTCCTTGACAAATTTGATCCG | Genotyping |
| *myc4*-RP | CGCTACACACACCATTGTTTG | Genotyping |
| *jin1-2-* FP | CAACGGGTAACGCGGCTTG | Genotyping |
| *jin1-2-* RP | CATCCCAAACACTCCTCC | Genotyping |
| GABI-LB8760BP | GGGCTACACTGAATTGGTAGCTC | Genotyping |
| **Primers used for cloning purposes** | | |
| MYC3-CDS-FP | CACCATGAACGGCAACATCATCAATCAACTTC | Gateway |
| MYC3-CDS-RP | TCAATAGTTTTCTCCGACTTTCGTCATC | Gateway |
| MYC4-CDS-FP | CACCATGTCTCCGACGAATGTTCAAGTAACCG | Gateway |
| MYC4-CDS-RP | TCATGGACATTCTCCAACTTTCTCCG | Gateway |
| COP1-FP | CACCATGGAAGAGATTTCGACGGATC | Gateway (BiFC) |
| COP1-RP | CGCAGCGAGTACCAGAACTTT GATG | Gateway (BiFC) |
| HY5-FP | CACCATGCAGGAACAAGCGACTAG | Gateway (BiFC) |
| HY5-RP | AAGGCTTGCATCAGCATTAGAACC | Gateway (BiFC) |
| **Primers used for gene expression analysis** | | |
| HY5-FP | CATCAAGCAGCGAGAGGTCA | qPCR |
| HY5-RP | CCGACAGCTTCTCCTCCAAA | qPCR |
| CAB1-FP | CTACCGACCCAGAGGCATTC | qPCR |
| CAB1-RP | AACTGGATCGGCCAAATGGT | qPCR |
| CHS-FP | CTCATGTCGTCTTCTGCACTACCT | qPCR |
| CHS-RP | GAAGCTTGGTGAGCTGGTAGTCA | qPCR |

| | | |
|---|---|---|
| CHI-FP | CCTTTTCGTCCTTGTTCTTCATCAT | qPCR |
| CHI-RP | GAGGCGGTTCTGGAATCTATCA | qPCR |
| ELIP2-FP | TCAACGGGAGACTAGCAATG | qPCR |
| ELIP2-RP | CCGTCAGAGATCTGAGCAAA | qPCR |
| RBCS1A- FP | AGAGTTAACTGCATGCAGGTGTG | qPCR |
| RBCS1A- RP | CCAACTCGAATTCAACACAAGG | qPCR |
| EF1α- FP | TACGCCCCAGTTCTCGATTG | qPCR |
| EF1α- RP | GGCTTGGTTGGGGTCATCTT | qPCR |
